# CRISPR-Mediated Linearization of IDLV Donor Enables Precise Targeted Integration in Human Hematopoietic Stem Cells

**DOI:** 10.64898/2026.06.15.732298

**Authors:** Giulia Scalisi, Aboud Sakkal, Laurie Lacombe, Francesco Sarnari, Marine Rouillon, Michela Rosiello, Alexandra Tachtsidi, Paola Galbiati, Guillaume Corre, Julie Oustelandt, Giulia Pavani, Marine Laurent, Mike Firth, Marie As, Marcello Maresca, Ivan Peyron, Peter J. Lenting, Anne Galy, Annarita Miccio, Mario Amendola

**Affiliations:** Genethon; 91000 Evry, France; Integrare Research Unit UMR_S951, Université Paris-Saclay, Univerisite Evry, Inserm, Genethon, Evry, France; Center for Cellular and Molecular Therapeutics, The Children’s Hospital of Philadelphia, Philadelphia, PA, USA; Data Sciences and Quantitative Biology, Discovery Sciences, R&D, AstraZeneca, Cambridge, UK; Museum National d’Histoire Naturelle, INSERM U1154, CNRS UMR7196, Paris, France; Genome Engineering, Discovery Sciences, R&D, AstraZeneca, Gothenburg, Sweden; UMR_S1176, Inserm, Univ. Paris-Sud, Université Paris-Saclay, Le Kremlin-Bicêtre, France; ART-TG, Accelerator of Technological Research in Genomic Therapy - Inserm US35, Evry, France; Imagine Institute, INSERM UMR1163, 75015 Paris, France; Department of Clinical and Experimental Medicine, University of Foggia, Foggia, Italy

## Abstract

*Ex vivo* genome editing of human hematopoietic stem and progenitor cells (HSPCs) requires targeted integration strategies that support large therapeutic DNA payloads while preserving stem cell fitness. Although CRISPR/Cas9-mediated homology-directed repair using AAV donors is effective, it is constrained by limited cargo capacity and adverse effects on long-term HSPCs function.

Integrase-defective lentiviral vectors (IDLVs) offer an alternative donor platform, yet their precise and controlled genomic integration remains inefficient.

Here, we describe TILV (Targeted Integration of Lentiviral Vector), a CRISPR-assisted knock-in strategy that exploits Cas9-mediated linearization of episomal IDLV DNA to expose a single homology arm and engage homology-mediated end-joining repair pathways.

TILV enables precise, directional and seamless integration of transgenes in multiple loci, enabling constitutive or physiological expression.

Using single-cell clonal analyses and targeted long-read sequencing, we define the molecular features of TILV-mediated integration and demonstrate preferential use of CRISPR-linearized episomal substrates. TILV supports accurate insertion of large therapeutic transgenes, without compromising HSPC viability or multilineage potential.

We further show that transient modulation of DNA repair pathway, in combination with extended homology arms, enhances integration efficiency and junctional precision.

Importantly, optimized TILV enables targeted integration in phenotypically defined long-term HSPCs, highlighting its potential for scalable and durable gene therapy.

## Introduction

Over the past decade, gene addition has emerged as a successful therapeutic strategy to engineer human hematopoietic stem and progenitor cells (HSPCs). Lentiviral vectors (LVs) are currently the gold standard for *ex vivo* HSPC gene addition and have been successfully applied to the treatment of multiple monogenic disorders, including primary immunodeficiencies, metabolic diseases, and haematological disorders (1–3). Stable genomic integration of LV-encoded transgenes enables long-term expression of functional proteins, supporting metabolic correction, clearance of toxic substrates, and cross-correction in affected tissues (4–6). However, LV integration occurs in a semi-random manner across the genome (7), raising safety concerns related to insertional mutagenesis, particularly when integrations disrupt tumour suppressor genes or occur near oncogenes and cis-regulatory elements (8–10). Moreover, transgene expression levels can vary widely among transduced cells, as it is influenced by both vector copy number and the local chromatin environment at the integration site (11–14).

To address these limitations, three main strategies have been developed: (i) engineering the viral integrase or its cellular cofactor LEDGF/p75 to bias LV integration toward selected chromatin regions (15,16); (ii) hybrid systems combining integrase-deficient lentiviral vectors (IDLVs) with transposons (e.g., Sleeping Beauty, PiggyBac) to exploit their intrinsic sequence preferences (17,18); (iii) targeted integration (TI) of donor DNA (dDNA) templates through homology-directed repair (HDR) mediated by site-specific nucleases, including CRISPR/Cas9, TALENs, and ZFNs, in combination with (19–21). Among these approaches, only HDR enables precise control over both the integration site and the number of integrated copies per cell.

Currently, the gold standard approach for targeted genome editing in HSPCs combines CRISPR/Cas9 nucleases with Adeno-Associated Virus serotype 6 (AAV6) donor template (22–26). Although very efficient *in vitro*, both AAV delivery and HDR based repair present important limitations. AAVs have a restricted packaging capacity (∼4.7 kb), which limits the size of the expression cassette. In addition, the p53-mediated DNA damage response (DDR) triggered by both Cas9 induced DSBs and AAV6 transduction can impair HSPCs proliferation (27), induce cellular senescence and negatively affect long-term engraftment potential (28). IDLV represents a valuable alternative donor-DNA delivery platform, although their use has remained limited by suboptimal integration efficiency.

IDLVs carry a catalytically inactive integrase that allows reverse transcription of the viral RNA genome into double-stranded DNA while preventing its genomic integration, resulting in episomal DNA forms. This feature makes IDLVs attractive dDNA templates, offering both a larger cargo capacity (∼10kb) and reduced cellular toxicity compared to AAV6 (27). Nevertheless, IDLV-based strategies generally show lower *ex vivo* and *in vivo* TI efficiencies than AAV6-mediated HDR (22,27). Beyond delivery constraints, the HDR repair pathway itself presents intrinsic biological limitations. HDR is predominantly active in proliferating cells, restricting its efficiency in long-term HSPCs, which are largely quiescent or slowly cycling (29). In addition, HDR requires dDNA to be flanked by two sequences, generally around 800 bp each, homologous to the genomic integration site (30,31). These homology arms can reduce vector packaging efficiency and limit the size of the deliverable expression cassette, owing to generating secondary structures, cryptic splice sites, polyadenylation signals, or GC-rich sequences (32–34). Furthermore, sequence mismatches introduced by single-nucleotide polymorphisms (SNPs; ∼1 per 1000 bp) within the homology arms can further compromise HDR-mediated integration efficiency (35,36).

To overcome the limitations of HDR, alternative TI strategies that exploits distinct DNA repair pathways have been developed. Homology-independent targeted integration (HITI) relies on non-homologous end joining (NHEJ) and employs dDNA cassette flanked by the same single-guide RNA (sgRNA) target sequences as the genomic site (37,38). This approach allows efficient knock-in in both dividing and non-dividing cells and has been described for AAV6 integration in HSPCs (39–41). However, HITI frequently introduces indels at vector-genome junctions (39), and requires repeated DSBs to achieve correct cassette orientation, thereby increasing genomic and cellular toxicity. Instead, in the PITCh (Precise Integration into Target Chromosome) system the plasmid dDNA cassette is flanked by short homology arms (5–25 bp) together with nuclease cleavage sites that allow donor linearization. PITCh therefore exploits the microhomology mediated end joining (MMEJ) pathway (42,43), that is active throughout most of the cell cycle, overcoming the S/G2 restriction associated with HDR (44,45). Finally, homology-mediated end joining (HMEJ) uses a similar dDNA cassette but incorporates longer homology arms (∼800 bp) (36,46,47). To date, however, these alternative end-joining-based strategies have not been evaluated and optimised for primary human HSPCs.

Here, we present ***T****argeted **I**ntegration of **L**entiviral **V**ector* (TILV), a novel CRISPR assisted strategy for dDNA TI into human HSPCs combining IDLV templates with CRISPR/Cas-based genome editing. By optimizing IDLV transduction, homology arm length, and pharmacological modulation of DNA repair pathway, we achieved efficient and precise integration of therapeutic transgenes in primary human HSPCs. We also characterized the integration mechanism using single-cell analysis and targeted long-read sequencing, establishing a versatile platform for precise gene addition with the potential to improve the safety profile of gene therapy approaches.

## Materials & Methods

### CRISPR/Cas9 nuclease

Recombinant *Streptococcus pyogenes* (Sp)Cas9 protein (with 2 nuclear localization signals, NLS, expressed in E. coli strain BL21 Rosetta 2) used in all experiments, was kindly provided by J.P. Concordet. Single guide RNAs (sgRNA) for *AAVS1* (5’-GTCCCCTCCACCCCACAGTG-3’) *HBA15 (*5’-GGGTTCTCTCTGAGTCTGTG-3’) and *HBA16* (5’-GTCGGCAGGAGACAGCACCA 3’) were previously described (25). RNP complexes were assembled by incubating (at 1:2 ratio) SpCas9 protein with synthetic sgRNA from Synthego for 10 min at RT with Alt-R Cas9 Electroporation Enhancer (Cat#1075915, Integrated DNA Technologies), unless otherwise indicated.

For targeted long-read sequencing, sgRNAs were designed as previously described for nanopore Cas9-targeted sequencing (nCATS) (48), selecting pairs of sgRNAs targeting opposite DNA strands and flanking the region of interest (ROI). The ROI encompassed the expected gene-editing cut sites within the *HBA1/HBA2* locus and spanned 14.3 kb. sgRNA design was performed using the CHOPCHOP online tool (49), prioritizing guides with the highest predicted on-target activity and minimal off-target potential. sgRNA was assembled as a duplex by annealing synthetic CRISPR RNAs (crRNAs; Integrated DNA Technologies, custom-designed) with trans-activating crRNAs (tracrRNAs; Cat# 1072532, Integrated DNA Technologies, custom-designed). The sequences of all sgRNAs used in this study are reported in Supplementary Table S1.

### IDLV donor templates

The design of the TILV transfer vectors for *AAVS1* and *HBA* editing is schematized in Figure 2a and Figure 3a. AAVS1 cargo cassette contains the GFP reporter under the human *PGK* promoter. and it is flanked at the 5’ by a single MH (35 bp long) and a sgT-RNA cut site (23 bp) for the sgRNA *AAVS1* (5’-GTCCCCTCCACCCCACAGTG GGG-3’). A cassette without MH (TILV-c Δ5’MH) was generated as control. HBA cargo cassette contains a promoterless GFP reporter flanked at the 5’ by a single MH (54 bp long) or LH (400 bp) and a sgT-RNA cut site (23 bp) for the sgRNA *HBA15* (5’-GGGTTCTCTCTGAGTCTGTGGGG −3’). Upon successful integration, transgene translation starts from the same ATG as the endogenous α-globin ATG for 5′ UTR integration [25-26]. A cassette without sgT-RNA target site (TILV-c Δsg-T) was generated as control. According to the experiments, *GFP* sequence was replaced with therapeutic cDNAs: *LAL* (Gene ID: 3988) cDNA with a C-terminal 3xHA tag (1xHA: TATCCCTATGACGTGCCTGATTACGCC) (50,51) and B domain deleted *F8* (V3: GCCACCAACGTGAGCAACAACAGCAACACCAGCAACGACAGCAACGTGAGC (52) R1654H (53) (Gene ID: 2157). Homology arm for 5′UTR AAVS1 integration is chr19:55115736:55115790 (hg38). Homology arm for 5′UTR α-globin integration is chr16:172642-172886 (hg38).

Plasmids and sequences are available from the corresponding author upon reasonable request and subject to a Material Transfer Agreement (MTA).

### IDLV production

IDLV were produced by transient transfection of HEK293T using third-generation lentiviral packaging plasmids (54). The transfer vector was co-transfected with pMDLg/p.RRE.D64V, pK.REV, and pseudotyped with the vesicular stomatitis virus glycoprotein G (VSV-G) envelope. IDLV were titrated in HCT116 cells and K562 cells. HIV-1 Gag p24 content was measured by ELISA (Cat# NEK050, Perkin-Elmer) according to manufacturer’s instructions. Vector titers and p24 measurements are reported in Supplementary Table S2.

### Cell lines and primary cells culture

K562 cells (ATCC® CCL-243) were cultured in RPMI 1640 medium supplemented with 2 mM GlutaMax (Cat# 35050-38, GIBCO^TM^), 10% fetal bovine serum (FBS, Cay#10270-106 GIBCO^TM^), 10 mM HEPES (Cat# H38871, Sigma Aldrich), 1 mM sodium pyruvate (Cat#11360-070 GIBCO^TM^) and 100 U/ml penicillin, 100 g/ml streptomycin (Cat#15140-122, GIBCO^TM^).

Human peripheral blood (PB) CD34^+^ cells were purified in house by immunomagnetic selection with EasySep™ Human Cord Blood CD34 Positive Selection Kit II (Cat#17896) from adult human peripheral blood provided by Etablissement Français du Sang (EFS), according to the agreement between Genethon and EFS (2023-2026-010-CCPSL GENETHON). G-CSF mobilized PB CD34^+^ HSPCs were purchased frozen by CliniSciences. HPSCs were seeded at the concentration of 5×10^5^ cells per ml in prestimulation media: serum-free StemSpan SFEM (Cat#09650, STEMCELL Technologies) supplemented with 100 U/ml penicillin, 100 g/ml streptomycin (Cat#15140-122, GIBCO^TM^), 2 mM GlutaMax (Cat# 35050038, GIBCO^TM^), 300 ng/ml hSCF (Cat#130-096-685, Milteny Biotech), 300 ng/ml hFlt3-L (Cat#130-096-479, Milteny Biotech), 100 ng/ml hTPO (Cat#130-095-752, Milteny Biotech), 20 ng/ml IL3 (Cat#130-095-069, Milteny Biotech), 750 nM and Stemregenin-1 (Cat#72342, STEMCELL Technologies) and 350 µM UM171 (Cat#72912, STEMCELL Technologies), 10 µM 16,16-Dimethyl Prostaglandin E2 (dmPGE2; Cat#14750, Cayman Chemical Company) and 4 µg/ml Protamine Sulfate (Cat# P3369, Sigma Aldrich). dmPGE₂ and protamine sulfate were added at the beginning of the culture, 2 h prior to IDLV transduction. All cells were cultured in a 5% CO2 humidified atmosphere at 37 °C.

### Gene editing K562

K562 cells (5 × 10^5^/ml) were transduced at multiplicity of infection (MOI) of 50 with the indicated IDLV in the figures, in cell culture medium containing 8 µg/ml Polybrene (Cat#TR-1003-G, Merck Millipore) as transduction enhancer (TE). Unless otherwise indicated, 24 hrs after transduction cells were washed in DPBS (Cat#10010023, Gibco™) and 2.5 × 10^5^ of transduced cells were electroporated using Nucleofector Amaxa 4D (Lonza, Basel, Switzerland) with SF Cell Line 4D-Nucleofector Kit (Cat# V4XC-2032, Lonza) (FF120 program). Cells were harvested 3-7-16-21 days after editing to analyze by flow cytometry the percentage of GFP positive cells and genomic (g)DNA was extracted for molecular analysis. Where indicated, K562 single clones were sorted after 22 days of editing by limiting dilution. For NHEJ inhibition experiment, 24 hrs after transduction cells were resuspended at the concentration of 0.5 x 10^6^ cells/ml and pre-incubated for 5 h with AZD7648 (Cat#S8843, Selleck Chemicals) at 1 µM prior to nucleofection. After electroporation, AZD7648 was added to the culture medium for an additional 24 hrs. Cells were then washed in DPBS and processed for analysis as described above.

### Gene editing HPSCs

Freshly purified or thawed mPB-derived HSPCs were cultured in prestimulation media for 48 h before the gene editing protocol, described in Figure 4a. Briefly, IDLV transduction of HSPCs was performed in retronectin-coated plates (5 µg/cm2; Cat# T100B, Takar). 3 × 10^5^ HSPCs (1 × 10^6^/ml) were seeded for 2 hour in prestimulation media plus 10 µM 16,16-Dimethyl Prostaglandin E2 (Cat#14750, Cayman Chemical Company) and 4 µg/ml Protamine Sulfate (Cat# P3369, Sigma Aldrich). Cells were transduced at MOI 250 with IDLV in presence of 100 µM LentiBoost (Cat# SB-A-LF-901-01, Sirion Biotech). After 24 hrs, cells were washed with DPBS and electroporated with RNP complex using P3 Primary Cell 4D-Nucleofector kit (Cat# V4XP-3032, Lonza) (CA137 program). In some experiments, IDVL transduction was performed with TE (Ciclosporin H, Cat# SML1575, Rapamycin Cat# R0395, Sigma Aldrich). Unless differently indicated, 1 day after editing, HSPCs were cultured in erythroid differentiation medium (serum-free StemSpan SFEM Cat#09650, STEMCELL Technologies, supplemented with 100 U/ml penicillin, 100 g/ml streptomycin Cat#15140-122, GIBCO^TM^, 2 mM GlutaMax Cat# 35050038, GIBCO^TM^, 300 ng/ml hSCF Cat#130-096-685, Milteny Biotech, 20 ng/ml IL3 Cat#130-095-069, Milteny Biotech, Epo 1 u/ml, IL3 5 ng/ml, Dexamethasone 2 µM Cat# D4902, and Betaestradiol 1 µM Cat# E2758 Sigma-Aldrich) For NHEJ inhibition experiment, cells were resuspended at the concentration of 0.5 x 10^6^ cells/ml in prestimulation media and pre-incubated for 5 h with AZD7648 (Cat#S8843, Selleck Chemicals) at 1 µM prior to nucleofection. DMSO was used as vehicle control. After nucleofection, AZD7648 was added to the culture medium for an additional 24 hrs before washout.

### Clonogenic assay

Colony-forming-unit cell assay was performed 1 day after editing, unless otherwise specified, by plating in two technical replicate 1000 HSPCs/each in methylcellulose-based medium (MethoCult H4435, STEMCELL Technologies) for 14 days. Colonies were counted and identified according to morphological criteria (BFU-E, CFU-G/GM, and CFU-GEMM). For figure 8e and figure 8m, BFU-E single colonies were manually picked and analyzed for vector copy number (VCN) quantification and In-out PCR at 5’vector-genome junction. Primer and probes sequences are listed in Supplementary Table S4. For figure 8h, bulk colonies were stained for CD45 and CD253a for immunophenotyping by FACS. Antibodies used are listed in Supplementary Table S3.

### Flow cytometry

Immunophenotypic analyses were performed on the fluorescence activated cell sorting (FACS) CytoFLEX S and LX (Beckman Coulter, Pasadena, CA, USA) For extracellular staining, cells (0.05-0.1 × 10^6^ cells) were harvested by centrifugation at 500 g for 5 minutes, followed by surface staining in a final volume of 100 µl, for 30 minutes at 4C in DPBS + 2% heat-inactivated FBS, in presence of Fc blocking reagent (Cat#564220, BD Pharminger). Samples were fixed by incubation in IOTest 3 1X (Beckman Coulter Cat# A07800).

For LAL-HA tag intracellular staining, cells were harvested at day 12 of erythroid differentiation, centrifuged at 500 g for 5 minutes and washed in FACS buffer. Surface antigens were stained in FACS buffer for 30 minutes at 4°C. Cells were washed once in FACS buffer, followed by a wash in cold DPBS. Cells were fixed using DPBS + 4% (v/v) PFA (Cat# 043368.9M, Alfa Aesar) for 20 minutes at room temperature. Next, cells were incubated with permeabilization buffer (Saponin 0.1% w/vol, 0.01% BSA in DPBS) for 5 minutes at room temperature. Intracellular HA-tag staining was performed in permeabilization buffer containing primary anti-HA TAG antibody (clone 16B12, Cat# 901514, BioLegend, 0.5 mg/sample) for 1 hour at 4°C. After washing with permeabilization buffer, samples were incubated with IgG1 PE anti-mouse, secondary antibody (Clone REA1017, REAfinity™, Cat #130-117-098, Milteny Biotech, 1:500 v/v) in permeabilization buffer for 30 minutes at 4°C and then washed twice with permeabilization buffer. Subsequently, FACS buffer + 1% (v/v) PFA was added to the samples and acquired with CytoFLEX S.

Cell sorting was performed with Beckman-Coulter - Astrios-EQ equipped with four lasers: blue (488 nm), yellow/green (561 nm), red (640 nm) and violet (405 nm). Cells were sorted with an 85 mm nozzle. Sheath fluid pressure was set at 45 psi. A highly pure sorting modality (four-way purity sorting) was chosen. Sorted cells were collected in 1.5 ml Eppendorf tubes containing 500 ml of DPBS or HSPC medium. Antibodies are listed in the Supplementary table S3. Data were analyzed with FCS Express 6 and FlowJo v10.x

### DNA analysis

Genomic DNA was extracted with QIAamp DNA Micro Kit (Cat#56304, Qiagen) or QuickExtract™ DNA Extraction Solution (Cat# QE09050 (Lucigen / Biosearch).

For indels quantification, 50 ng or 5μL of gDNA were used to amplify the region that spans the cutting site of each sgRNA using KAPA2G Fast ReadyMix (Cat# 07960891001, Kapa Biosystem). After Sanger sequencing the percentage of indels was calculated using Inference of CRISPR Edits (ICE) analysis with Synthego Performance Analysis software (version 3.0; Synthego, Redwood City, CA, USA; 2019) (55) or Tide software (56). For single clone analysis, 5μL of gDNA were used for In-out PCR on vector-genome 5’ and 3’junctions using KAPA2G Fast ReadyMix.

Droplet digital PCR (ddPCR)was performed according to manufacturer’s instruction using ddPCR Supermix for Probes No dUTP (Cat#1863023 Biorad) and 1–50 ng of genomic DNA digested with HindIII (Cat# R0104S, New England Biolabs). Droplets were generated using AutoDG Droplet Generator and analyzed with QX200 droplet reader, according to the manufacturer’s instructions. Data analysis was performed with QuantaSoft (Biorad, Hercules, CA, US). To quantify on-target transgene integration events, primers and probe were designed spanning the 5’vector-genome junction. Human albumin (ALB) was used as reference for copy number evaluation Primer and probes sequences are listed in Supplementary Table S4.

### Gene expression analysis

Total RNA was purified using RNeasy Micro kit (Cat# 74004 Qiagen), according to the manufacturer’s instructions and DNase treatment was performed using RNase-free DNase Set (QIAGEN). Complementary (d)DNA was reverse-transcribed using Transcriptor First Strand cDNA Synthesis Kit (Cat# 0489703000, Roche). qPCR was performed using Maxima Syber Green/Rox (Cat# K0222, Life Scientific, Thermo-Fisher Scientific) on LightCycler 480 PCR system (Roche). The relative expression of each target gene was normalized using human GAPDH as a reference gene (NM_002046.6) and represented as 2^ΔCt for each sample or as fold changes (2^ΔΔCt) relative to the control. Primer sequences are listed in Supplementary Table S4.

### Protein analysis

We quantified FVIII antigen in supernatant with AlphaLISA human FVIII detection kit (Cat# AL342HV, Revvity) according to the manufacturer’s instructions Briefly, supernatants were collected 12 days after editing and 10 ul were incubated with 5X AlphaLISA Anti-hFactor VIII Acceptor beads 10 µg/mL for 30 minutes at 23°C. 20 µL of 5X Biotinylated Anti-hFVIII Antibody (1 nM) was added and samples were incubated 60 minutes at 23°C. 25 µL of 2X SA-Donor beads (40 µg/mL final) was added and samples were incubated 30 minutes at 23°C in the dark. Luminescence was read at 615 nm on EnSpire Multimode Plate Reader (PerkinElmer). Data were calculated by plotting the AlphaLISA standard curve versus the concentration of analyte, to a nonlinear regression using the 4-parameter logistic equation.

FVIII activity was measured using a chromogenic assay using the Biophen FVIII-assay kit (Cat#227102 Hyphen BioMed) as instructed. Human normal plasma was used as a reference. LAL enzymatic activity was detected in undiluted or 1:2 diluted supernatants, as previously described (50). Briefly, samples were incubated 10 min at 37 °C with 150 µM Lalistat-2 (Cat# SML2053, Sigma-Aldric), a specific competitive inhibitor of LAL, or water. Samples were then transferred to a Optiplate 96 F plate (Cat# 6005270, PerkinElmer) where fluorimetric reactions were initiated with 75 μl of substrate buffer, (340 μM 4-MUP (4-methylumbelliferyl palmitate, Cat#16089, Cayman Chemical Company), 0.9% Triton X-100 (Cat# T8787, Sigma-Aldric) and 220 µM cardiolipin (Cat#C-0563, Sigma-Aldric) in 135 mM acetate buffer pH 4.0). After 10 min, fluorescence was recorded (35 cycles, 30” intervals, 37 °C) using SPARK TECAN Reader (Tecan, Austria). Kinetic parameters (average rate) were calculated using Magellan Software. LAL activity over untreated samples was quantified using this formula:

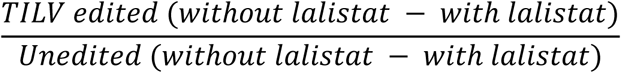

### Amplicon sequencing analysis

InDel quantification from human genomic DNA was performed by Amplicon-Sequencing. Briefly, genomic DNA (gDNA) from 10^5^ cells was extracted using the QuickExtract™ DNA Extraction Solution (Cat# QE09050, Lucigen). For the TILV characterization with short homology arm, a target-specific amplification was performed with Phusion™ High-Fidelity DNA Polymerase (Cat# M0530S, New England Biolabs) to enrich the target region and include specific adapters. The amplicons were gel-purified using NucleoSpin Gel and PCR clean-up (Cat# 740609, Macherey Nagel) following the manufacturer’s instructions. Next, the libraries were amplified by PCR using barcoded primers and Phusion™ High-Fidelity DNA Polymerase, followed by gel-purification.

For the characterization of LH-TILV, a selective first PCR was performed to exclude episomal IDLV DNA. PCR was carried out using one primer annealing to the genomic HBA locus and one primer annealing to GFP, thereby selectively amplifying HBA integrated TILV. The resulting PCR product was then used as template for the second PCR reaction to enrich the target region and include specific adapters.

Primers are listed in Supplementary table S4. Amplicon libraries were sequenced using Illumina technology on the NovaSeq platform in paired-end mode (2 × 151 bp) (Macrogen). After base calling, the generated BCL files were converted into FASTQ files using bcl2fastq software. Genome editing variant analysis was performed using CRISPResso2 (v2.2.12) (57), run with the parameters and reference sequences provided in Supplementary Table S5. To further analyze and visualize indel profiles, we used a Python implementation of RIMA (58), which takes as input the CRISPResso2 output, using a quantification window set to 1.

### Targeted long read sequencing

Cells were collected on day 28 after editing for K562 and around 10-12 days post erythroid differentiation for HPSCs. At least 4 x10 ^6^ cells were collected. High molecular weight DNA was extracted from K562 or HSPCs with the Nanobind CBB kit (Cat# 102-301-900, PacBio) according to manufacturer’s instructions. DNA was size selected using Short Read Eliminator XS kit (Cat# 102-208-200, PacBio) and quantified using the Qubit 4 fluorometer (Cat# Q33226, Thermo Fisher Scientific). 5µg of genomic DNA was subjected to Cas9-mediated PCR-free enrichment protocol (version: CAS_9106_v109_revC_16Sep2020) before library preparation using the Cas9 Sequencing Kit (Cat# SQK-CS9109, Oxford Nanopore Technologies-ONT). Libraries were loaded onto MinION R9.4.1 flow cells (Cat# FLO-MIN106D, Oxford Nanopore Technologies-ONT,) and run on GridION using MinKNOW software.

We selected reads longer than 4kb using cutadapt (v4.9) and removed adaptor sequences using porechop (v0.2.4). Trimmed reads were aligned to the reference genome (GRCh38) using minimap2 (v2.28). Reads overlapping the enrichment region were selected and realigned on the locus regions with or without the *HBA* deletion using optimized parameters to account for the high degree of homology between *HBA2* and *HBA1* genes. Variants calling programs Sniffle (v1.0.12) and CuteSv (v1.0.8) were used to identified indels and complex genomic rearrangements (DUP, INV, TRANS). Samtools (v1.21) was used to collect locus coverage and pileup alignments. Split alignments were annotated using the plasmid and locus features. Bed files were then post-processed using the R program to reconstruct reads structure from split alignments annotations a-d produce tables and figures.

### GUIDE seq

DNA library preparation for GUIDE-seq analysis was performed as previously described (59). K562 were nucleofected with 80 pmol of dsODN and a Cas9/sgRNA RNP (1:2 ratio). After 5 days in culture, DNA was isolated using the QIAamp DNA mini kit following manufacturer’s instructions (Cat# 51304, QIAGEN). DNA was sonicated (Bioruptor Pico sonicator) to generate fragments of 400–900 bp. These fragments were ligated to adaptors, and two steps of DNA amplification were performed using the KAPA HyperPrep Kit (Cat# KK8504, Roche). The libraries were then purified, and concentration was measured using the qPCR KAPA libraries quantification kit (Cat# 07960140001, Roche) and Qubit device (Thermo-Fisher Scientific) Average length was estimated with the Tape Station bioanalyzer 2100 (Agilent). The libraries were pooled, diluted to a final concentration of 4nM and loaded onto a MiSeq flow cell (Institute Imagine Genomics Facility)Subsequent steps included demultiplexing, consolidation of PCR duplicates, identification of on-target cleavage sites, detection of off-target activity, and data visualization, which were performed using an GENETH’OFF GUIDE-Seq Analysis pipeline (<u>GitHub - gcorre/GENETHOFF: Versatile GuideSeq analysis pipeline · GitHub</u>) (60).

## Results

### TILV enables precise integration into a safe harbor locus

To create TILV, we designed an IDLV donor in which the expression cassette is positioned downstream of a single homology arm and a single guide RNA target site (sgT-RNA), referred to as TILV-cassette (TILV-c). Following nuclear entry, the IDLV genome is reverse-transcribed into double-stranded DNA, which can persist in the nucleus either as linear DNA or, more commonly, as episomal forms (61), predominantly 2 LTR circles generated through NHEJ or 1 LTR circles generated by HDR mediated LTR–LTR recombination (62) (**Fig. 1a–b**). Upon Cas9/sgRNA cleavage, the circular IDLV donor is linearized exposing the homology arm for recombination with the genomic DSB (**Fig. 1c**).

**Figure 1.**
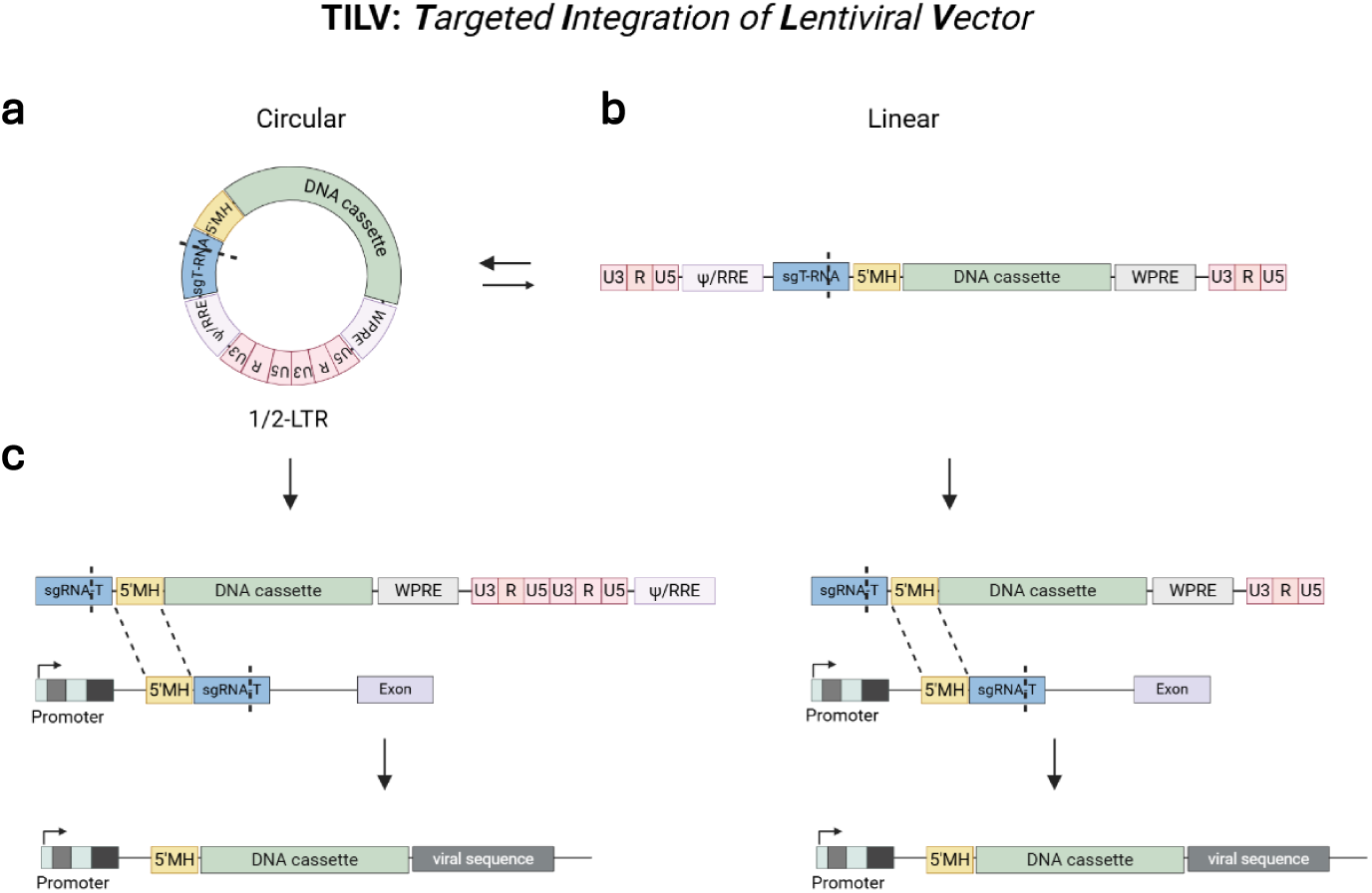
Schematic representation of TILV system. The TILV system is based on IDLV encoding a TILV cassette containing a transgene preceded by a 5’ microhomology arm (5’MH) and a sgRNA target site (sgT-RNA). Following transduction, the IDLV can persist in either (**a**) a circular episome or, (**b**) less frequently, in a linear form. (**c**) Upon delivery of Cas9 and the corresponding sgRNA, both the genomic target site and the sgT-RNA sequence within the TILV cassette are cleaved. This cleavage event linearizes the donor and exposes the microhomology region, thereby facilitating directional transgene integration into the genome through microhomology-mediated DNA repair pathways. Created with BioRender.com

To test TILV-mediated TI, we selected the AAVS1 safe-harbor locus in the human K562 erythroleukemia cell line (25). We generated a TILV donor encoding green fluorescent protein (GFP) under the control of the human phosphoglycerate kinase (PGK) promoter, preceded by a 55-bp 5′ microhomology arm (5′MH) and the sgRNA-T site (**Fig. 2a**). To evaluate the contribution of the MH to the TI, we generated a control donor vector lacking the MH sequence (TILV-c Δ5’MH) (**Fig. 2a**).

**Figure 2.**
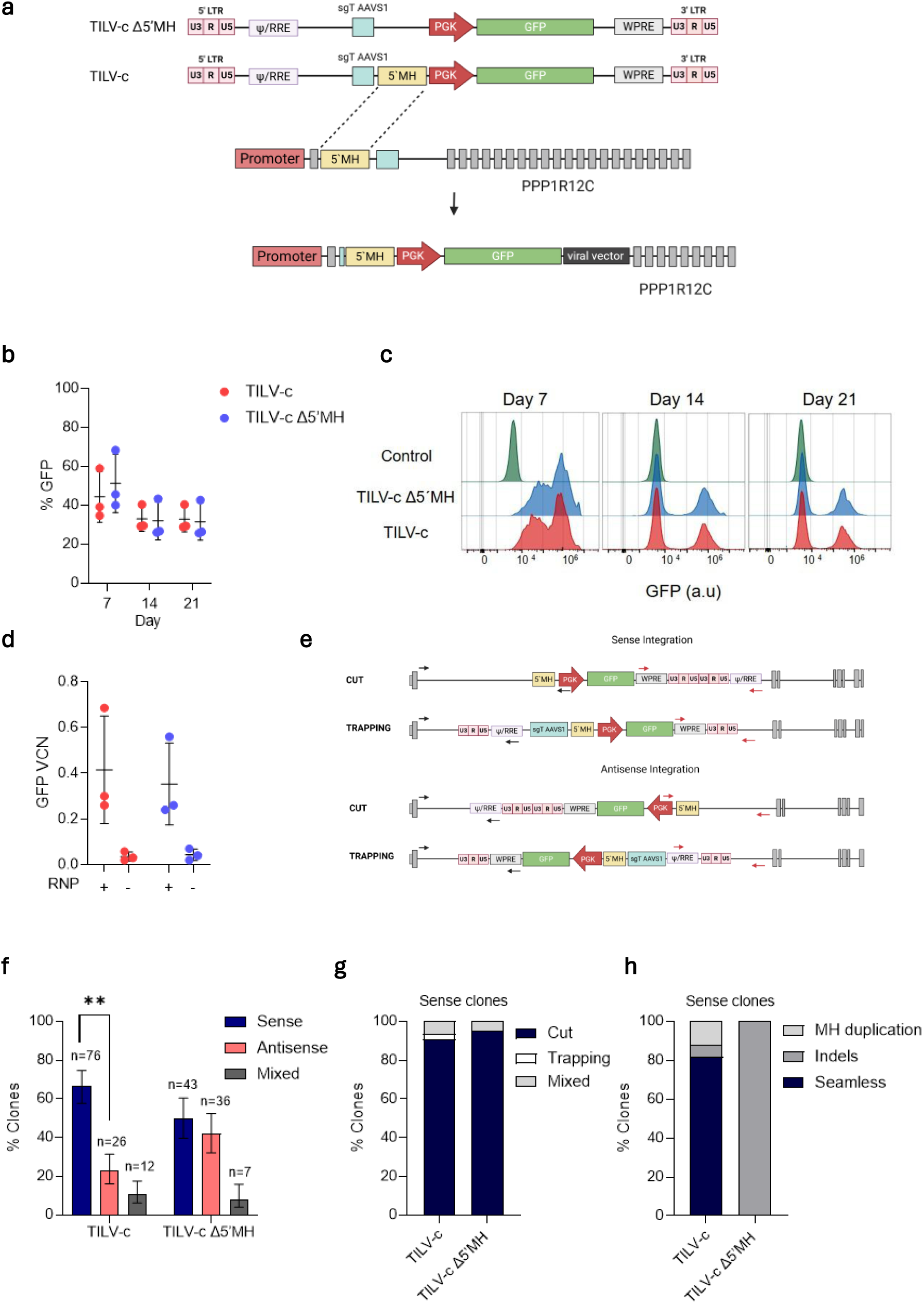
TILV-c cassette enables precise integration into the AAVS1 safe harbor locus. **a**) Design of the TILV-c targeting the *AAVS1* locus. The donor encodes the GFP (728 bp) with the human phosphoglycerate (PGK) promoter downstream of a 5’microhomology (MH) and the sgRNA target site (sgT-AAVS1). A control cassette lacking 5’MH (TILV-c Δ5’MH) is used to assess the contribution of microhomology to TI. Created with BioRender.com. **b-c**) Percentage (b) and expression level (c) of GFP-positive K562 cells at day 7, 14 and 21 following editing with TILV-c or TILV-c Δ5’MH (n=3, mean ± SD). **d**) Vector copy number (VCN) of GFP in K562 edited with TILV-c and TILV-c Δ5’MH at day 14 with (+) or without (-) RNP (n=3, mean ± SD). **e**) Schematic representation of the in–out PCR strategies used to analyze dDNA orientation and integration mechanism (primer sequences are listed in Supplementary Table S4). Red and black arrows indicate the primer positions relative to the genome and to TILV-c: black primers amplify upstream and red primers downstream regions of the transgene. Created with BioRender.com. **f**) Edited K562 clones were analyzed for sense or antisense TILV integration. Clones positive for both PCRs were classified as mixed. TILV-c (n=138) and TILV-c Δ5’MH (n=114) clones were compared by Pearson’s Chi-square test (fractions ± 95% CI; error bars indicate the upper and lower limits, ** p=0.01). **g**) Classification of integration mechanisms among clones with sense integrations: “Cut” (donor cleaved prior to integration, n=69 TILV-c; n=40 TILV-c Δ5’MH), “Trapping” (uncut donor captured n=2 TILV-c), or “Mixed” (both mechanisms detected in the same clone, n=5 TILV-c; n=2 TILV-c Δ5’MH). **h**) PCR amplifications of vector-genome 5’ junctions of each clone (g) were Sanger sequenced to assess integration fidelity. Junctions were classified as seamless (n = 27 TILV-c), indel-containing (n = 2 TILV-c; n = 22 TILV-c Δ5′MH), or exhibiting MH duplication (n = 4 TILV-c).

K562 cells were transduced with each of the two vectors and, 24 hrs later, electroporated with Cas9/sgRNA ribonuclear complex (RNP). At 7, 14, and 21 days post-editing, we checked sgRNA indels efficiency (**Supplementary Fig. S1a**) and GFP expression by flow cytometry. Both donor constructs resulted in comparable levels of GFP-positive cells over time (**Fig. 2b–c, Supplementary Fig. S1b**). This observation was further confirmed by droplet digital PCR (ddPCR)-based quantification of GFP copy number (**Fig. 2d**), indicating similar integration efficiency between the two donor designs.

To assess whether the presence of a MH arm influences the orientation and precision of TI, we isolated single-cell GFP positive clones (185 and 181 for TILV-c and TILV-c Δ5’MH, respectively). First, we confirmed that each clone stably expressed GFP at similar level. (**Supplementary Fig. S1c–d**).

Then, we performed genomic PCR analysis to determine: (i) the orientation of cassette integration (sense or antisense); (ii) the Cas9-mediated TILV cleavage (linearised vs trapping integration, respectively); (iii) the precision of the 5’genome–vector junction (**Fig. 2e**).

After excluding PCR negative clones (47 and 67 TILV-c and TILV-c Δ5′MH clones, respectively), integration analysis revealed a clear bias toward TILV sense integration, with 55% (76/138) of TILV-c clones showing sense-orientation integration against 18% (26/138) with anti-sense one. In contrast, TILV-c Δ5’MH clones showed a near-random orientation distribution, with similar proportion of sense (38%, 43/114) and antisense (32%, 36/114) integrations (**Fig. 2f**).

The remaining clones displayed multiple donor insertions, with 1 on-target in sense integration (**Fig. 2f**).

Interestingly, both donor constructs were mostly integrated after Cas9 linearization, as evidenced by genome–vector junction analyses (**Fig. 2g, Supplementary Fig. S1e**).

In addition, sequence analysis of the 5′ genome–vector junctions revealed that TILV-mediated integration was predominantly seamless, consistent with MH-dependent engagement of MMEJ (**Fig. 2h**). In contrast, TILV-c Δ5’MH clones frequently exhibited indels (**Fig. 2h, Supplementary Fig. S1f**), indicative of error-prone NHEJ-mediated integration.

Finally, we examined whether donor DNA design reduce off target integration. Using CRISPOR (63) and COSMID (64) webtools we identified the top 8 predicted sgRNA off-targets (OTs) (**Supplementary Fig. S1g**), four of which were confirmed by GUIDE-Seq (59,60) (**Supplementary Fig. S1h**). By PCR analysis, we validated off-target 8 (**Supplementary Fig. S1i**) and we quantified donor DNA integration events at this site. Interestingly, TILV-c exhibited a lower frequency of unintended integration events compared to TILV-c Δ5’MH donor (17% versus 25%, respectively) (**Supplementary Fig. S1j**).

Overall, these results demonstrate that the presence of the MH within the TILV donor enables directional and precise on-target integration of the DNA cassette, while limiting nonspecific donor trapping at off-target sites.

### TILV allows hijacking the HBA endogenous promoter for transgene expression

We next investigated whether this strategy could be applied to achieve physiologically regulated transgene expression under the control of an endogenous human promoter. We selected the human α-globin (*HBA*) locus as its promoter displays robust and erythroid-specific transcriptional activity, making it a interesting platform for therapeutic gene addition in HSPCs (25,26,65).

We designed a promoterless TILV-c carrying a GFP reporter downstream of a 56-bp MH and an sgRNA target site, enabling TI immediately downstream of the endogenous *HBA* promoter (**Fig. 3a**).

**Figure 3.**
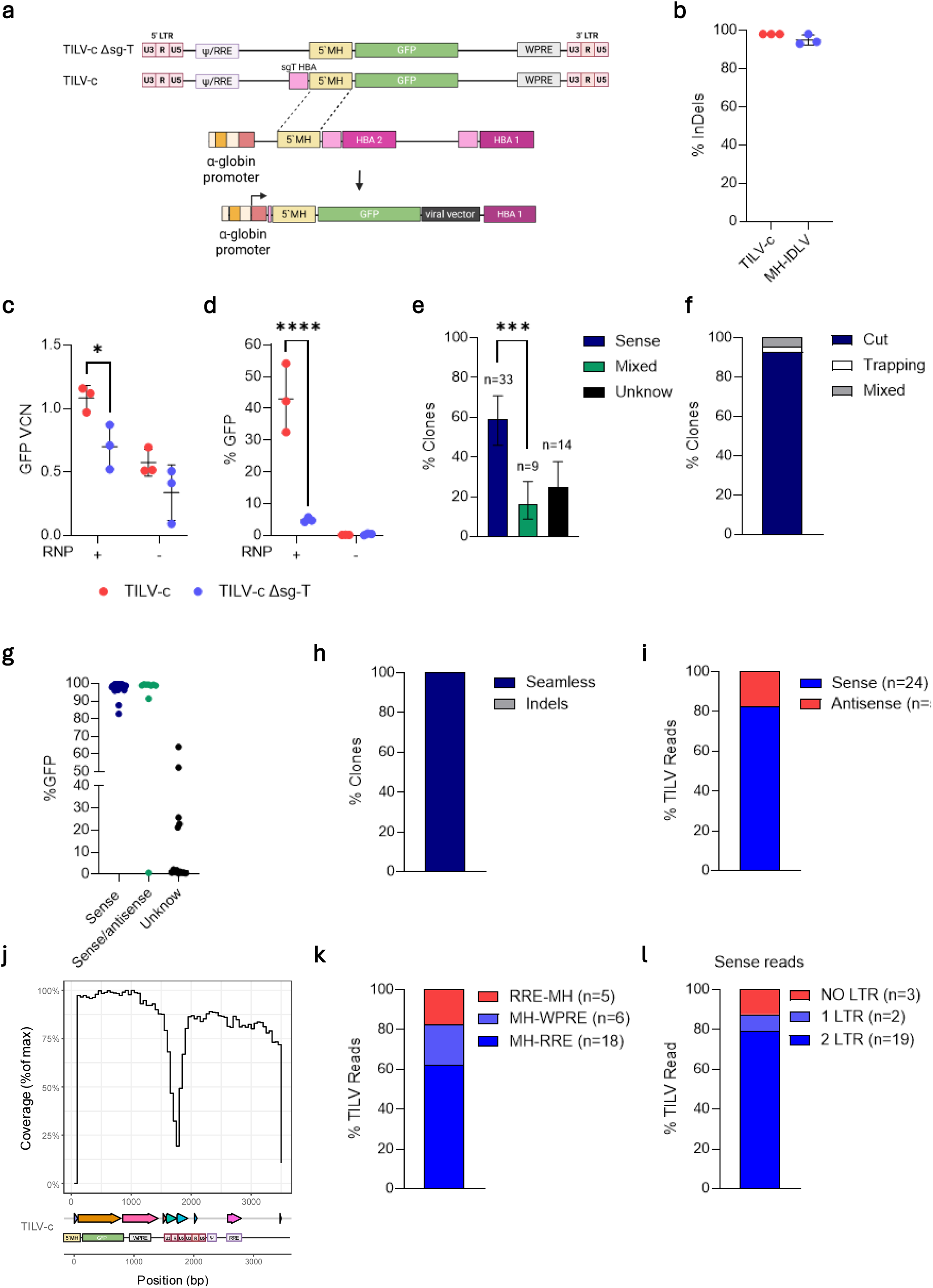
TILV system enables precise transgene expression driven by the endogenous HBA promoter. **a**) Design of the TILV cassette targeting the *HBA* locus. *GFP* is downstream of the 5’MH and the sgRNA-T (sgT-HBA). The control TILV-c Δsg-T lacks the sgT-HBA site. Created with BioRender.com. **b**) Editing efficiency in K562 cells 7 days after editing, expressed as percentage of indels at the target site (n=3, mean ± SD). **c**) VCN of GFP in K562 edited with TILV-c or TILV-c Δsg-T at day 14 post editing (n=3, mean ± SD; two-way Anova test, *p=0.01). **d**) Percentage of GFP-positive K562 cells on day 14 after editing, edited (+) or transduced-only (-) (n=3, mean ± SD; two-way Anova test, ***p<0.0001). **e**). A total of 178 K562 clones were isolated by serial dilution and analyzed by in-out PCR for sense or antisense TILV-c integration. Clones positive for both PCRs were classified as mixed, while PCR-negative clones were classified as unknown (n=56, Chi-squared test, fraction ± 95% CI, ***p=0.0001). **f**) Classification of integration mechanisms among sense and sense/antisense single-cell clones: “Cut” (donor cleaved prior to integration, n=39), “Trapping” (uncut donor captured, n=1), or “Mixed” (both forms detected in the same clone, n=2). **g**) Percentage of GFP-positive cell clones show in (e). **h**) PCR amplifications of 5’ vector-genome junctions of each clone were Sanger sequenced to assess the integration fidelity (n=33). **i**) tLRS analysis of cassette orientation in TILV-edited K562. **j**) tLRS read coverage across 4 kb region of chromosome 16 spanning the *HBA* locus (n=2 biological replicate). Below are indicated the different TILV-c elements: MH (microhomology), GFP (green fluorescent protein), HIV-1-5’ LRT/HIV-1-3’ LRT (long terminal repeat), Ψ (Psi), WPRE (Woodchuck Hepatitis Virus (WHV) Posttranscriptional Regulatory Element), RRE (Rev Response Element). **k**) Analysis of 3’ vector-genome junctions. Integration orientations were assigned as follows: RRE-MH (from RRE to MH, antisense), MH–WPRE (from MH to WPRE), and MH–RRE (from MH to RRE). **l**) Number of LTR sequences detected within each tLRS read.

We first defined the optimal timing between TILV transduction and RNP delivery. Although on-target cleavage efficiencies were comparable across all tested conditions (**Supplementary Fig. S2a**), both GFP integration and expression increased progressively when RNP was transfected 4–24 hrs after TILV transduction, reaching a plateau between 24 and 48 h (**Supplementary Fig. S2b–c**). Based on these findings, we selected 24 hrs post-transduction as the optimal time points for RNP electroporation in subsequent experiments.

To test whether donor linearization is required for efficient TI at the *HBA* locus, we compared the TILV-c donor with a control vector lacking the sgRNA target site (TILV-c Δsg-T; **Fig. 3a**). K562 cells were transduced with either TILV and electroporated with RNP. While genomic cleavage efficiencies and transduction levels were similar between conditions (**Fig 3b; Supplementary Fig. S2d**), TILV-c yielded higher GFP copy number and expression after 14 days (**Fig. 3c–d**). These data indicate that Cas9-mediated donor linearization facilitates TI at the *HBA* locus. Noteworthy, TILV integration was obtained also when Cas9/sgRNA was delivered as plasmid, highlighting the flexibility of the strategy with respect to nuclease delivery modalities (**Supplementary Fig. S2e–g**). To further validate these findings, we exploited two distinct sgRNAs: one (*sgHBA15*) targeting both the genome and the donor cassette, and a second (*sgHBA16*) targeting only the genome (**Supplementary Fig. S2h**). K562 cells were transduced with TILV and, 24 hrs after, transfected with different RNPs. Despite comparable genomic cleavage between the two guides (**Supplementary Fig. S2i**), GFP integration and expression were detected only in cells edited with *sgHBA15* (**Supplementary Fig S2j–k)**. This result confirms that simultaneous cleavage of the donor molecule is necessary to enable productive TIat the endogenous HBA promoter.

Together, these experiments demonstrate that efficient TILV-mediated targeting of the HBA locus requires Cas9-dependent donor linearization and is maximized when RNP delivery is performed 24 h after IDLV transduction.

### Quantitative and structural characterization of TILV-mediated transgene integration

To evaluate the accuracy of TILV-c integration into the HBA locus, we isolated 178 edited K562 single-cell clones. Among these, we selected 56 clones with at least 1 integrated GFP copy, of which 40% possessed a VCN ≥2 (**Supplementary Fig. S3a**). PCR analyses (**Supplementary Fig. S3b**) revealed that 75% (42/56) of GFP positive clones harbored at least one sense integration, of which 16% (9/56) showed also antisense integration (**Fig. 3e**). This result was in line with flow cytometry analysis, showing that ∼99% of sense and mixed clones were positive for GFP (**Fig. 3g**). The higher number of mixed clones is probably due to the presence of 4 HBA per cells. In addition, we confirmed that TILV integrated following CRISPR-linearization of episomal IDLV (**Fig. 3f**). Importantly, sequence analysis of the 5′ genome–vector junctions showed that sense-oriented integrations were largely seamless, displaying precise junctions consistent with homology-mediated repair (**Fig. 3h**).

To obtain an unbiased and comprehensive profile of TILV-c integration events, we performed targeted long-read sequencing (tLRS) of the edited *HBA* locus (**Supplementary Fig. S3c**) on bulk TILV-c edited K562 cells. Approximately 16% of the reads contained an integrated cassette (**Supplementary Fig. S3d**), predominantly in sense-orientation (**Fig. 3i**), of which approximately 80% spanned the entire IDLV (from MH to RRE) and virtually 100% encompassed the full transgene expression cassettes (**Fig. 3j–k**). In addition, 75% of integrants retained two LTRs (**Fig. 3l**), suggesting that 2 LTR circular IDLVs serve as the primary substrate for TILV genomic integration.

We also detected tandem integrations of two or four cassette copies in 21% of cassette-positive reads (**Supplementary Fig. S3e**), likely arising from LTR-LTR recombination at episomal level. Collectively, these results demonstrate that the TILV system enables efficient, directional and precise transgene integration into the *HBA* locus. The integrated sequence is predominantly in the form of a monomeric, full-length expression cassette derived from the CRISPR-based linearization of a 2 LTR IDLV episomes.

### TILV enables targeted integration of therapeutic transgenes into primary human HSPCs

We next evaluated whether the TILV platform could support TI in primary human HSPCs, a clinically relevant cell type characterized by limited proliferative activity and heightened sensitivity to DNA damage. First, we optimized IDLV transduction conditions of HSPCs from healthy donors comparing commonly used transduction enhancers-dmPGE2 (16,16-dimethyl PGE2) (66,67), LentiBOOST (68,69), Cyclosporin H (70), and Rapamycin (71). After 24 hours, transduced HSPCs were transfected with *HBA* targeting RNP. While on-target cleavage efficiencies were comparable across all tested conditions (**Supplementary Fig. S4a),** we observed variable levels of GFP integration and expression (**Supplementary Fig. S4b-c**). Notably, no adverse effects on HSPCs viability and multilineage differentiation potential were observed, as confirmed by colony-forming cell (CFC) assays (**Supplementary Fig. S4d**). Based on these results, we selected the combination of ddPGE2 and LentiBOOST as the standard transduction protocol for subsequent experiments.

We then investigated the impact of Cas9-mediated donor linearization on integration efficiency in primary HSPCs (**Fig. 4a**) by comparing TILV-c and TILV-c Δsg-T control (**Fig. 3a**). Despite comparable genomic cleavage efficiencies (**Supplementary Fig. S4e**) and vector transduction (**Fig. 4b**), the TILV platform yielded higher GFP expression levels (**Fig. 4c**), without major alteration in clonogenic output or multilineage differentiation potential (**Fig. 4d–e**). To assess integration precision, we performed next-generation sequencing (NGS) of the 5′ genome–vector junctions, revealing that TILV-mediated integration in HSPCs is predominantly seamless, closely mirroring the high-precision profiles observed in K562 cells (**Fig. 4f**). We then performed Rational InDel Meta-Analysis (58) on indel-containing reads carrying the integrated cassette and observed a balanced distribution of small insertions and deletions due to alternative end-joining (alt-EJ) DNA repair pathways (**Supplementary Fig. S4f–g**), confirming the involvement of those DNA repair pathways in TILV-c integration.

**Figure 4.**
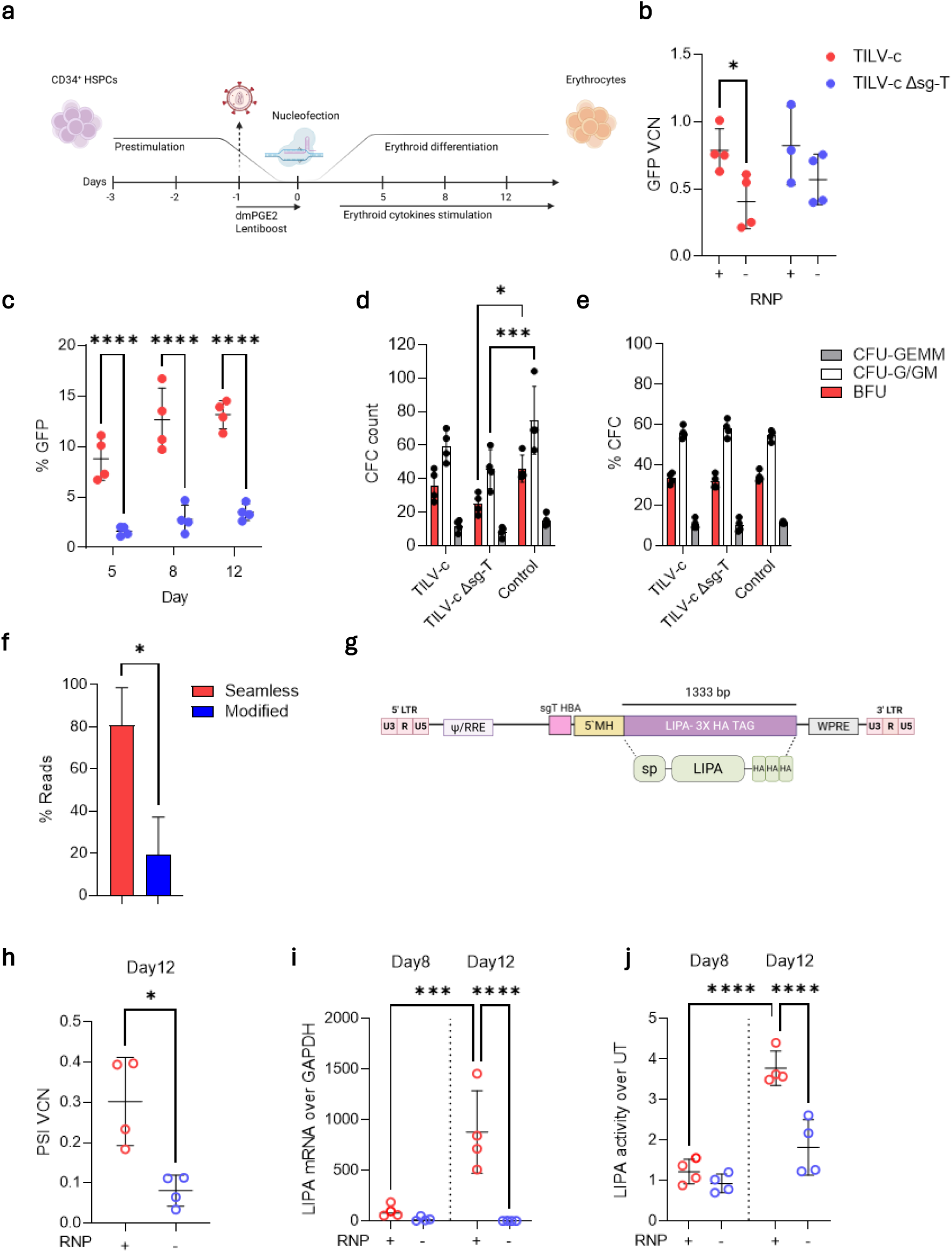
TILV enables efficient targeted integration in human HSPCs. **a**) Experimental workflow for HSPCs editing: prestimulation, IDLV transduction, Cas9/sgRNA nucleofection and *in vitro* erythroid differentiation. Created with BioRender.com. **b**) VCN of GFP in edited HSPCs with TILV-c or TILV-C Δsg-T after 12 days of erythroid differentiation, in edited (+) or transduced-only (-) HSPCs (n=3-4, mean ± SD; two-way Anova test, *p<0.05). **c**) Percentage of GFP-positive cells on day 5, 8 and 12 of erythroid differentiation in TILV-c and TILV-C Δsg-T edited HSPCs (n=4; two-way Anova test, ****p<0.0001). **d-e**) Colony-forming cell (CFC) number (d) and colony types (e) in edited HSPCs. CFU-GEMM: granulocyte, erythroid, macrophage, megakaryocyte; BFU-E: burst-forming unit-erythroid; CFU-G/GM: granulocyte-macrophage count (n=3, mean ± SD two-way Anova test, *p=0.01,***p<0.001). **f**) Amplicon sequencing of 5’ vector–genome junctions of TILV-c edited HSPCs after 12 days of erythroid differentiation (n=3, mean ± SD; unpaired t test, *p<0.05). **g**) Design of the TILV cassette coding for the *LIPA* cDNA flagged with 3 HA-TAG sequences, downstream of an optimized signal peptide. Created with BioRender.com. **h**) VCN of PSI (Ψ) as proxy for LIPA TILV integration, after 12 days of erythroid differentiation (n=4, mean ± SD; paired t test, *p<0.05). **i**) *LIPA* mRNA expression in edited HSPCs at days 8 and 12 of erythroid differentiation (n=4, mean ± SD; two-way Anova test, ***p<0.001, ****p<0.0001). **j**) Fold change of LAL enzymatic activity relative on untreated HSPCs (n=4, mean ± SD; two-way Anova test, ****p<0.0001).

We then tested the potential of the TILV system to the delivery of therapeutic transgenes with distinct sizes and clinical relevance. First, we inserted the lysosomal acid lipase transgene (LIPA; **Fig. 4g**), which is mutated in lysosomal acid lipase deficiency (OMIM #278000), a severe metabolic disorder characterized by lipid accumulation and multi-organ failure.

HSPCs were LIPA-TILV transduced and RNP HBA transfected (**Supplementary Fig. S4h**), resulting in integration (**Fig. 4h, Supplementary Fig. S4i)** and substantial LIPA mRNA and activity increase upon erythroid differentiation (**Fig. 4i–j**).

We then evaluated TILV-mediated integration of the B-domain-deleted form of Factor VIII (BDDF8, 4.4 kb) (52,53) (**Fig. 5a**), whose deficiency causes the hemophilia A coagulation disorder (OMIM #306700). Noteworthy, this cDNA is approximately six times larger than GFP and approaches the maximum packaging capacity of AAV vectors, even before accounting for the homology arms required for classical HDR mediated integration.

**Figure 5.**
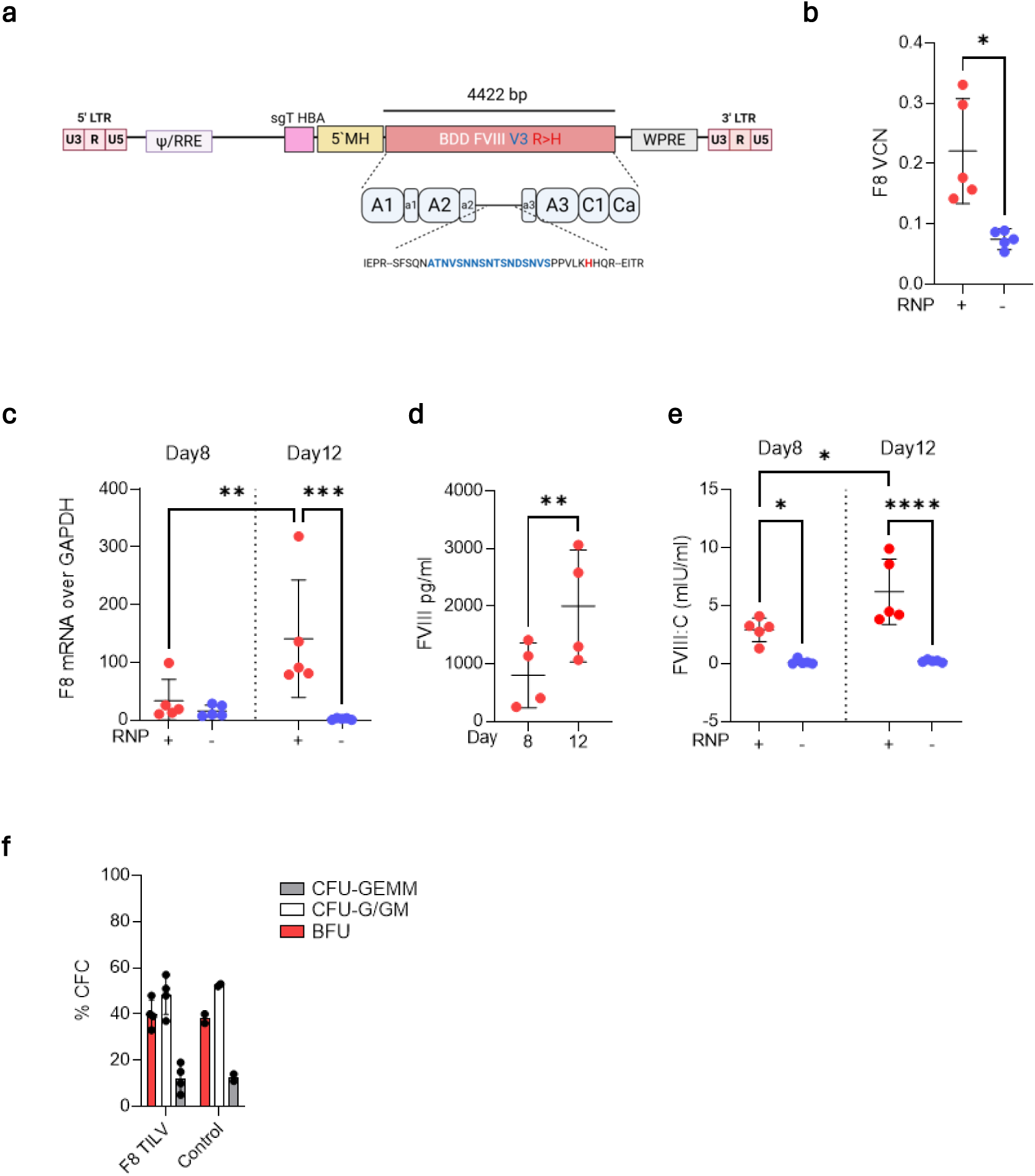
TILV system enables TI of a large therapeutic payload in HSPCs. **a**) Design of the TILV cassette coding for an optimized codon B domain deleted *F8* cDNA, with a 31-amino acid linker (52) and a R1645H (53) point mutation to increase protein secretion. Created with BioRender.com. **b**) VCN of the *F8* in F8 TILV after 12 days of erythroid differentiation (n=5, mean ± SD; paired t test *p<0.01). **c**) *F8* mRNA expression in edited HSPCs at days 8 and 12 of erythroid differentiation (n=5, mean ± SD, two-way Anova test, **p<0.01, ***p<0.001). **d**) Secreted FVIII protein level measured by AlphaLISA in the cell supernatant at day 8 and 12 days of erythroid differentiation (n=5, mean ± SD; paired t test, **p<0.01). **e**) FVIII functional activity in supernatants collected at day 8 and 12 of erythroid differentiation (n=5, mean ± SD; two-way Anova test, *p< 0.05, ****p<0.0001). **f**) CFC frequency in F8 TILV–edited HSPCs (n=2, mean ± SD).

Efficient FVIII integration and expression were observed both in K562 cells (**Supplementary Fig. S5a-e)** and HSPCs (**Supplementary Fig. S5f–g, Fig. 5b-e**). Importantly, the integration of this large genetic payload did not impair the multilineage differentiation potential of HSPCs (**Fig. 5f**), although a moderate reduction in clonogenic potential was observed (**Supplementary Fig. S5h**), as previously reported (27).

Taken together, these data demonstrate that TILV enables TI and expression of different therapeutic transgenes of varying sizes in primary human HSPCs.

### DNA-PK inhibition enhances TILV integration precision

Since the efficiency of TILV integration expression in HSPCs remained low, we investigated whether pharmacological modulation of DNA repair pathway choice could further enhance TILV performance.

Based on previous studies (72–77), we hypothesized that NHEJ inhibition could promote alternative DNA repair pathways, including MMEJ, and consequently TILV TI. We tested this hypothesis in K562 and human HSPCs transduced with a GFP encoding TILV and treated with AZD7648 DNA-PK inhibitor. As expected, AZD7648 treatment slightly reduced overall indels frequency and changed indels pattern, with an increase in junctional complexity (**Supplementary Fig. S6a–b**). Notably, the treatment almost doubled the number of GFP expressing cells (**Supplementary Fig. S6c)**. Since we did not observe a corresponding increase in TI efficiency (**Supplementary Fig. S6d**), this effect is most consistent with an increased frequency of sense-oriented and seamless integration events.

We next evaluated the effect of AZD7648 drug also on HSPCs, that exhibit distinct activation of DNA repair pathways compared to K562 cells (78,79) (**Fig. 6a**). In HSPCs, AZD7648 treatment led to a marked reduction in indels formation (**Fig. 6b**), with a significantly increased GFP expression during erythroid differentiation (**Fig. 6c–d**), without affecting cell viability (**Supplementary Fig. S6e)**. Similarly to K562, we did not observe any significant difference in GFP DNA copies (**Fig. 6e, Supplementary Fig. S6f**), indicating that enhanced transgene expression was not due to increased integration frequency. Instead, NGS of the 5′ genome-vector junction confirmed a substantial increase in integration precision, with fewer than 10% of junctions exhibiting indels in AZD7648-treated cells (**Fig. 6f**). Importantly, the indel spectrum of reads containing the integrated cassette was not markedly altered (**Supplementary Fig. S6 g–h**). Finally, AZD7648 treatment did not alter HSPC multilineage differentiation potential (**Fig. 6g**) or the erythroid lineage specific transcription of the integrated GFP (**Supplementary Fig. S6j**), despite a modest reduction in CFC capacity (**Supplementary Fig. S6i**).

**Figure 6.**
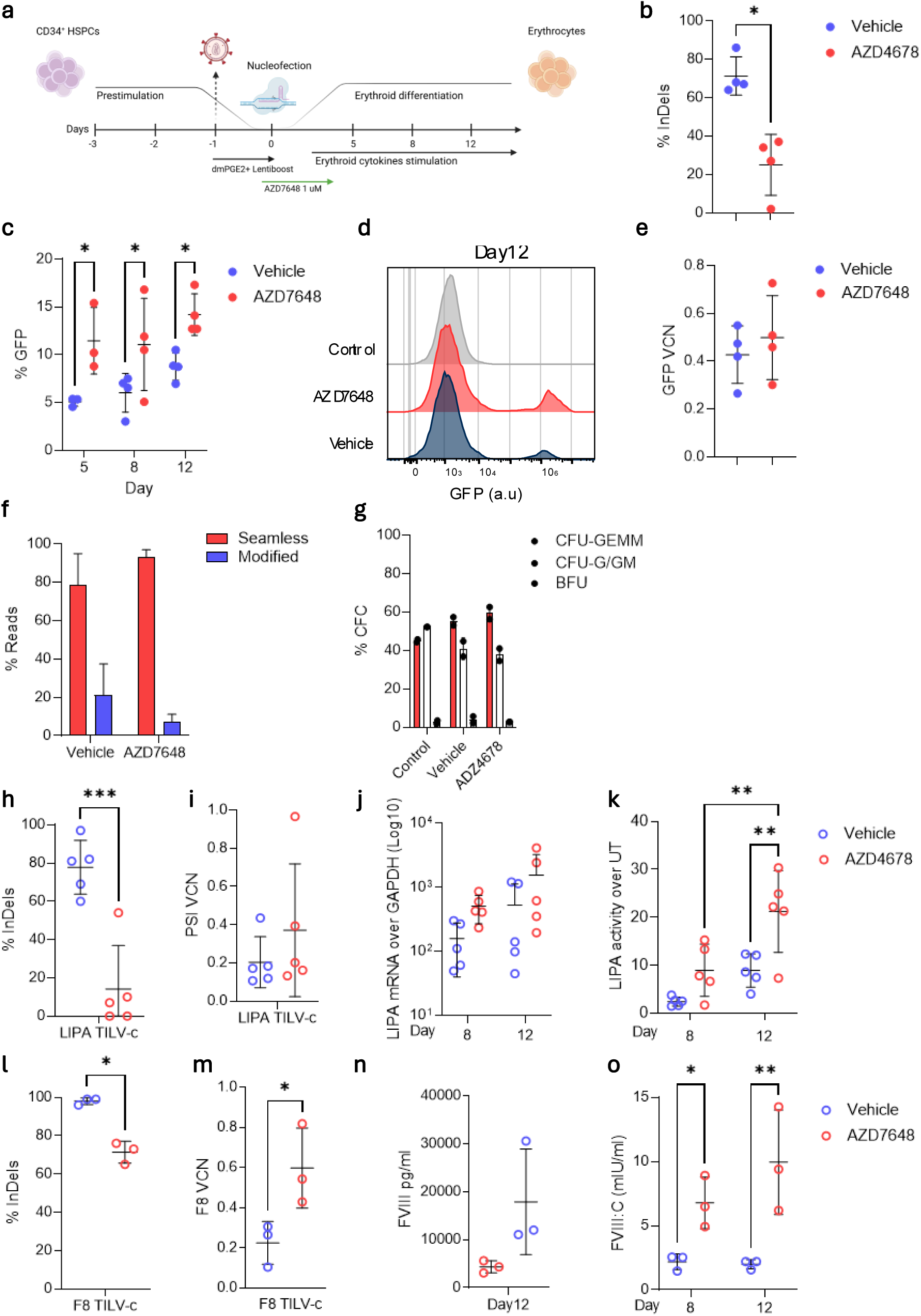
DNA-PK inhibition enhances TILV mediated TI in HSPCs. **a**) Timeline of HSPCs editing protocol using 1 µM AZD7648. Created with BioRender.com. **b**) Editing efficiency in HSPCs, at day 8 of erythroid differentiation expressed as percentage of indels (n=4, mean ± SD; paired t test, *p<0.05). **c-d**) TI efficiency measured as (c) percentage of GFP-positive cells at day 5, 8 and 12, and (d) GFP expression histogram at day 12 of erythroid differentiation in TILV edited HSPCs ± 1 µM AZD7648 (n=3, mean ± SD; two-way Anova test, *p<0.05). **e**) VCN of GFP in TILV edited HSPCs ± 1 µM AZD7648, at 12 days of erythroid differentiation. (n=4, mean ± SD; paired t test, ns). **f**) Targeted amplicon sequencing of 5′ vector–genome junctions in TILV edited HSPCs ± AZD7648, at 12 days of erythroid differentiation (n=3, mean ± SD; two-way Anova test, ns). **g**) CFC frequency in edited HSPCs (n=2, mean ± SD). **h**) Editing efficiency in HSPCs treated with MH-TILV LIPA cassette, at day 8 of erythroid differentiation, expressed as percentage of indels (n=5, mean ± SD; paired t test, ***p<0.001). **i**) VCN of PSI (Ψ), proxy for LIPA TILV, at 12 days of erythroid differentiation (n=5, mean ± SD; paired t test, ns). **j**) *LIPA* mRNA expression at days 8 and 12 of erythroid differentiation (n=5, mean ± SD, expressed in Log10 scale, two-way Anova test, ns). **k**) Fold change of LIPA enzymatic activity relative to untreated HSPCs (n=5, mean ± SD; two-way Anova test, **p<0.01). **l**) Editing efficiency in HSPCs treated with MH-TILV F8 cassette, at day 8 of erythroid differentiation, expressed as percentage of indels (n=3, mean ± SD; paired t test, *p<0.05). **m**) VCN of the *F8* in HSPCs treated with F8 TILV at 12 days of erythroid differentiation (n=3, mean ± SD, paired t test, p<0.05). **n-o**) FVIII protein protein levels (n) and functional activity (o) in the supernatant of F8 TILV edited HSPCs ± 1 µM AZD7648 at day 8 and 12 of erythroid differentiation (n=3, mean ± SD; paired t test, ns (n) or two-way Anova test, *p<0.05, **p<0.01).

The beneficial effects of DNA-PK inhibition were confirmed using the *LIPA* (**Fig. 6h–k**, **Supplementary Fig. S6 k–l)** and the *F8* (**Fig. 6 l–o**) TILVs, which both showed enhanced transgene expression and preserved cellular fitness.

Overall, these findings demonstrate that DNA-PK inhibition enhances both the precision of TILV-mediated TI and the efficiency of transgene expression in primary human HSPCs.

### Long Homology arm enhances TILV integration efficiency and precision in HSPCs

We then hypothesized that extending homology arm length could stabilize its interaction with the target genomic locus and thereby enhance TI efficiency in HSPCs. Thus, we replaced the 56 bp microhomology (MH) with a 400 bp long homology arm (LH) to generate the LH-TILV-c (**Fig. 7a**), aiming to leverage the homology-mediated end-joining (HMEJ) pathway (36,46,47)

**Figure 7.**
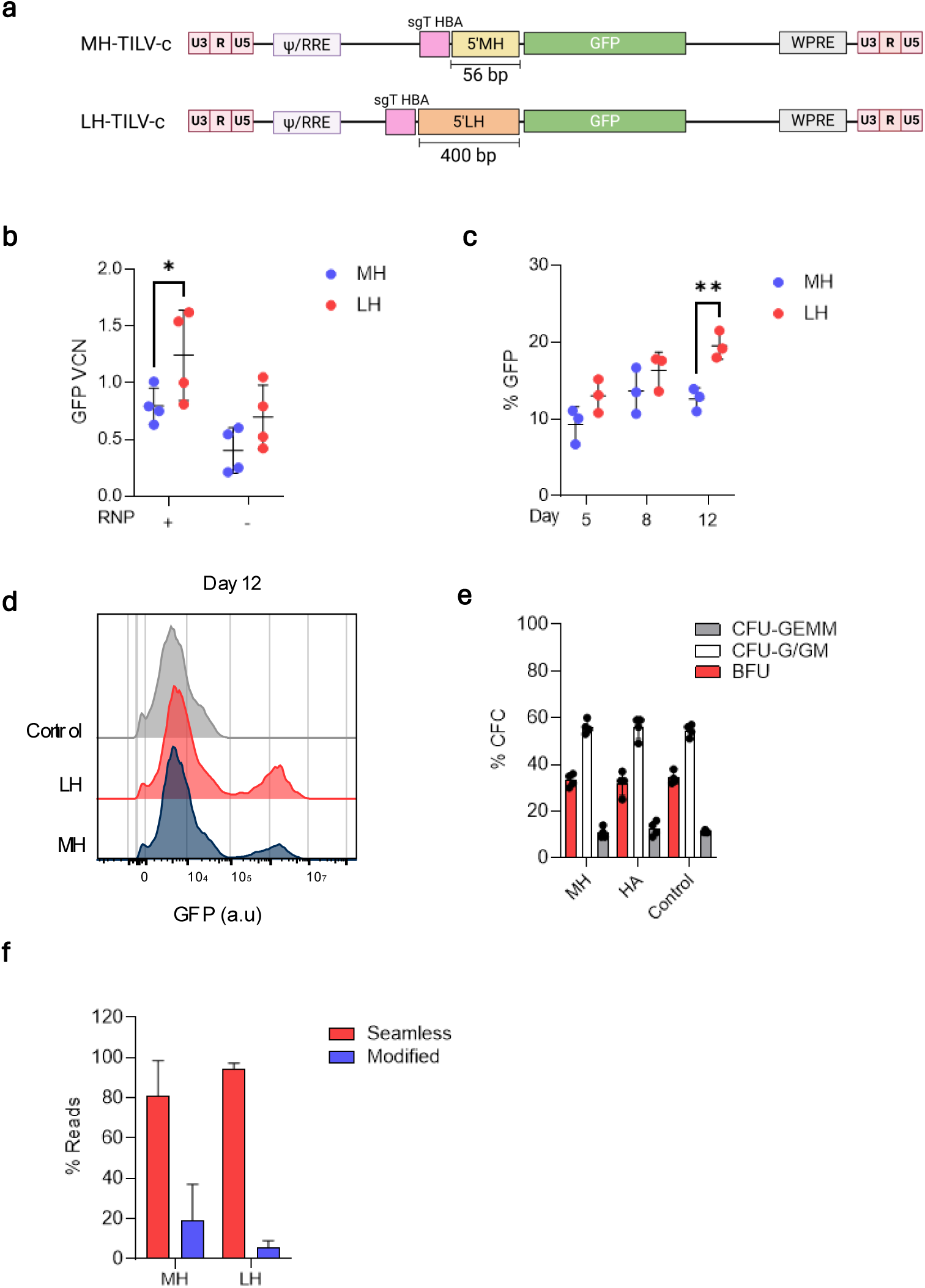
Long homology arms promote accurate and efficient genome editing in HSPCs. **a)** Design of the LH TILV cassette coding for GFP downstream a long single homology arm (400 bp). compared to the shorter homology arm (56 bp) of the TILV cassette (MH-TILV). Created with BioRender.com. **b**) VCN of GFP for MH and LH-TILV cassette at 12 days of erythroid differentiation, edited (+) or transduced-only (-) HSPCs (n=4, mean ± SD; two-way Anova test, *p<0.05). **c-d**) TI efficiency expressed as percentage of GFP-positive cells at day 5, 8 and 12, (c) and GFP expression histogram at day 12 of erythroid differentiation of MH and LH-TILV (d) (n=3-4, mean ± SD; two-way Anova test, **p<0.01). **e**) CFC frequency in edited HSPCs (n=2, mean ± SD). **f**) Targeted amplicon sequencing of 5′ vector–genome junctions from MH- and LH-TILV edited HSPCs after 12 days of erythroid differentiation (n=3, mean ± SD; two-way Anova test, ns).

In HSPCs, LH-TILV-c showed a clear increase in both GFP integration and expression compared to MH-TILV-c (**Fig. 7 b–d; Supplementary Fig. S7d**), without impairing HSPC multilineage differentiation potential (**Fig. 7e, Supplementary Fig. S7e**). NGS analysis of the 5’ genome-vector junction confirmed that LH-TILV-c further improved integration precision (**Fig. 7f**). Notably, the overall indel profile of reads carrying the integrated cassette remained largely unchanged (**Supplementary Fig. 7 Sf–g**). To a lower extend, similar results were obtained also in K562 (**Supplementary Fig. S7 a–c**).

Together, these data indicate that extending homology arm length significantly enhances both the efficiency and precision of TILV-mediated TI in human HSPCs, consistent with preferential engagement of HMEJ-like repair pathways.

### Combining DNA-PKi and LH improves targeted integration in HSPCs

Finally, we asked whether combining donor design optimization and DNA repair pathway modulation could further enhance TILV performance in primary human HSPCs. Specifically, we evaluated the effect of DNA-PK inhibition on LH-TILV-c–mediated integration, reasoning that canonical NHEJ may still compete with HMEJ even in the presence of long homology arms.

In HSPCs edited with LH-TILV-c, the AZD7648 treatment induced a marked increase in both GFP copies and expression (**Fig. 8 a–c, Supplementary Fig. S8a**), while maintaining high integration precision at the 5′ genome–vector junction (**Fig. 8d**, **Supplementary Fig. S8b–c**). Consistent with previous observations (25), GFP expression was restricted to erythroid differentiated CFC cells, faithfully recapitulating the endogenous pattern of HBA promoter activity **(Fig. 8 e-f**). By analyzing 52 single BFUE, we observed that 40% of CFC were GFP positive, with ∼60% of these harboring a single, predominantly seamless integration event (**Fig. 8 g-h, Supplementary Fig. 8 Sd–e**).

**Figure 8.**
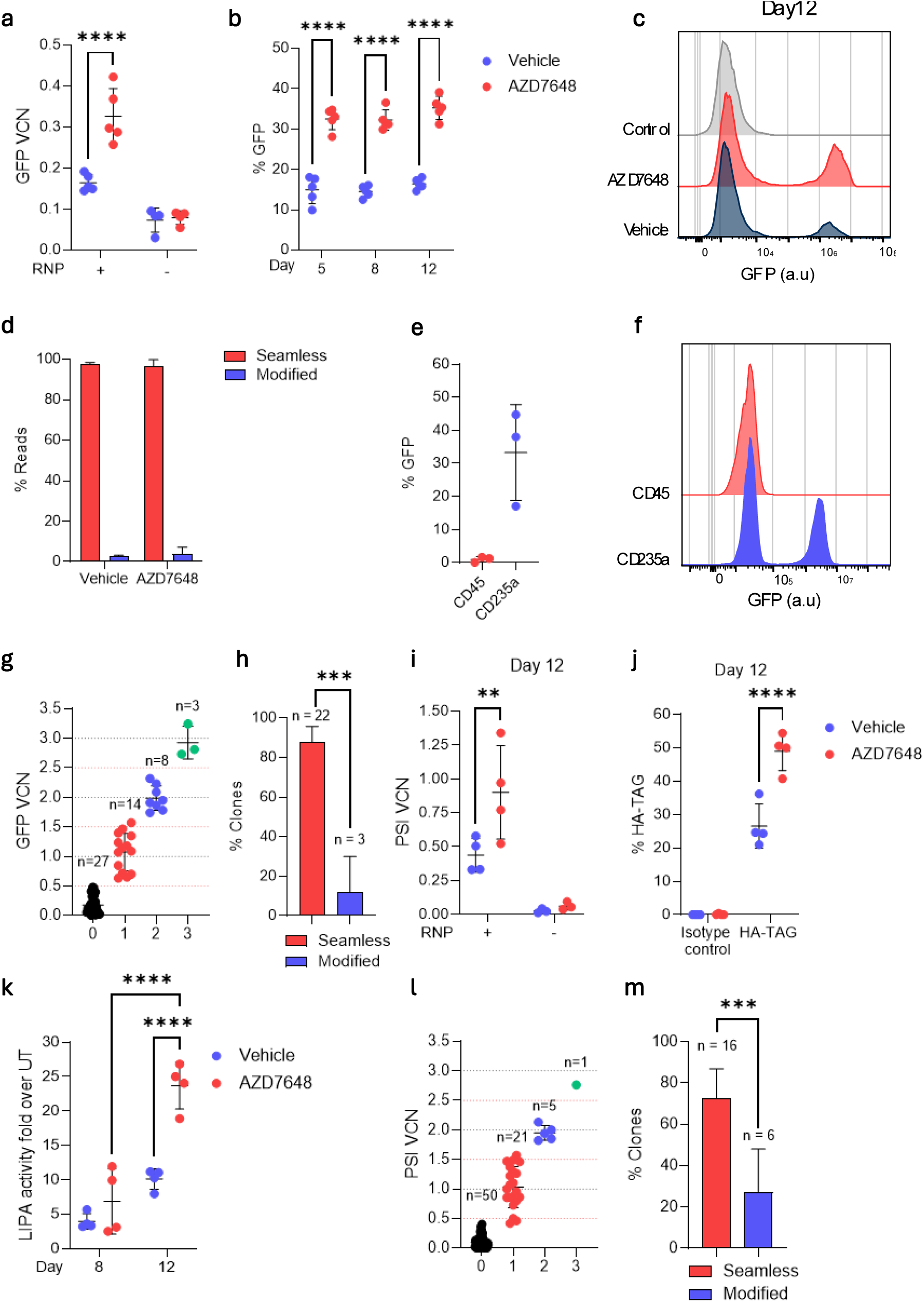
Combined use of LH TILV-c and NHEJ inhibition enable precise and lineage-specific expression of therapeutic transgenes. **a)** VCN of GFP for LH-TILV-c ± AZD7648 1 µM, at 12 days of erythroid differentiation, in edited (+) or transduced-only (-) HSPCs (n=4-5, mean ± SD; two-way Anova test, ***p<0.0001). **b-c**) TI efficiency expressed as (b) percentage of GFP-positive cells at day 5, 8 and 12, and (c) GFP expression histogram at day 12 of erythroid differentiation of LH TILV-c edited HSPCs ± AZD7648 1 µM (n=5, mean ± SD; two-way Anova test, ****p<0.0001). **d**) Targeted amplicon sequencing of 5′ vector–genome junctions in LH-TILV-c edited HSPCs ± AZD7648 1 µM, at 12 days of erythroid differentiation (n=3, mean ± SD, two-way Anova test, ns). **e-f**) TI efficiency in CFC as (**e**) percentage of GFP positive and (**f**) histogram of GFP expression of non-erythroid (CD45+, red plot) and erythroid (GYPA+, blue plot) CFC (n=3, mean ± SD; paired t-test, ns). **g**) VCN on single CFC of LH TILV-c edited HSPCs ± AZD7648 1 µM. Clones were classified as unmodified (VCN 0–0.5, black), monoallelic (VCN 0.5–1.5, red), or biallelic (VCN 1.5–3, blue/green) TILV integration. Each point represents an individual clone (n=52). **h**) Characterization of single BFU CFC clones by Sanger sequencing of 5′ genome–vector junctions (n=25; Exact Binomial test, error bars indicate the upper and lower limits, ***p<0.001). (**i**) VCN of PSI (Ψ), proxy for LIPA LH TILV-c, of edited HSPCs ± AZD7648 1 µM at 12 days of erythroid differentiation, edited (+) or transduced-only (-)(n=4, mean ± SD; two-way Anova test, *p<0.05). **j)** Percentage of HA-TAG positive (LIPA expressing) cells at 12 days of erythroid differentiation in LH-TILV-c edited HSPCs ± AZD7648 (n=4, mean ± SD; two-way Anova test, ****p < 0,0001). **k)** Fold change in LIPA enzymatic activity relative to untreated HSPCs (n=4, mean ± SD, two-way Anova test; ***p<0.001 and ****p < 0.0001). **l**) VCN in single CFC clones edited with LIPA LH-TILV-c ± AZD7648. Clones were classified as unmodified (VCN 0–0.5, black), monoallelic (VCN 0.5–1.5, red), or biallelic (VCN 1.5–3, blue/green) LH-TILV integration. Each point represents an individual clone (n=77). **m**) Characterization of single BFU CFC clones by Sanger sequencing of 5′ genome–vector junctions (n=22; Exact Binomial Test, error bars indicate the upper and lower limits, ***p<0.001).

These results were confirmed using the *LIPA* therapeutic transgene, where we achieved high levels of both integration and expression, with over 50% of transgene positive cells (**Fig.8 i–k; Supplementary Fig. S8f–h**). Around 35% of BFUE CFC colonies were positive for LIPA (**Supplementary Fig. S8i**), with nearly 80% carrying a single, mostly seamless integration (**Fig 8l-m, Supplementary Fig. S8j**).

To obtain a complete picture of the integration outcomes, we performed tLRS of LH-TILV-c–edited HSPCs. Approximately 23% of full-length on-target reads contained the integrated LH TILV-c (**Supplementary Fig. S9a**). Interestingly, almost 100% of integrations were monomeric (**Supplementary Fig. 9b**), with sense orientation (**Fig. 9a**) and spanning the entire expression cassette (from HA to WPRE; **Supplementary Fig. S3d, Fig 9 b–c**). Furthermore, we observed that ∼60% of the integrations originated from cleaved 2 LTR circular IDLVs, while the remaining events were consistent with integration of linear IDLV intermediates, terminating at the WPRE or LTR sequences (**Supplementary Fig. S3d, Fig 9 b–d**).

**Figure 9.**
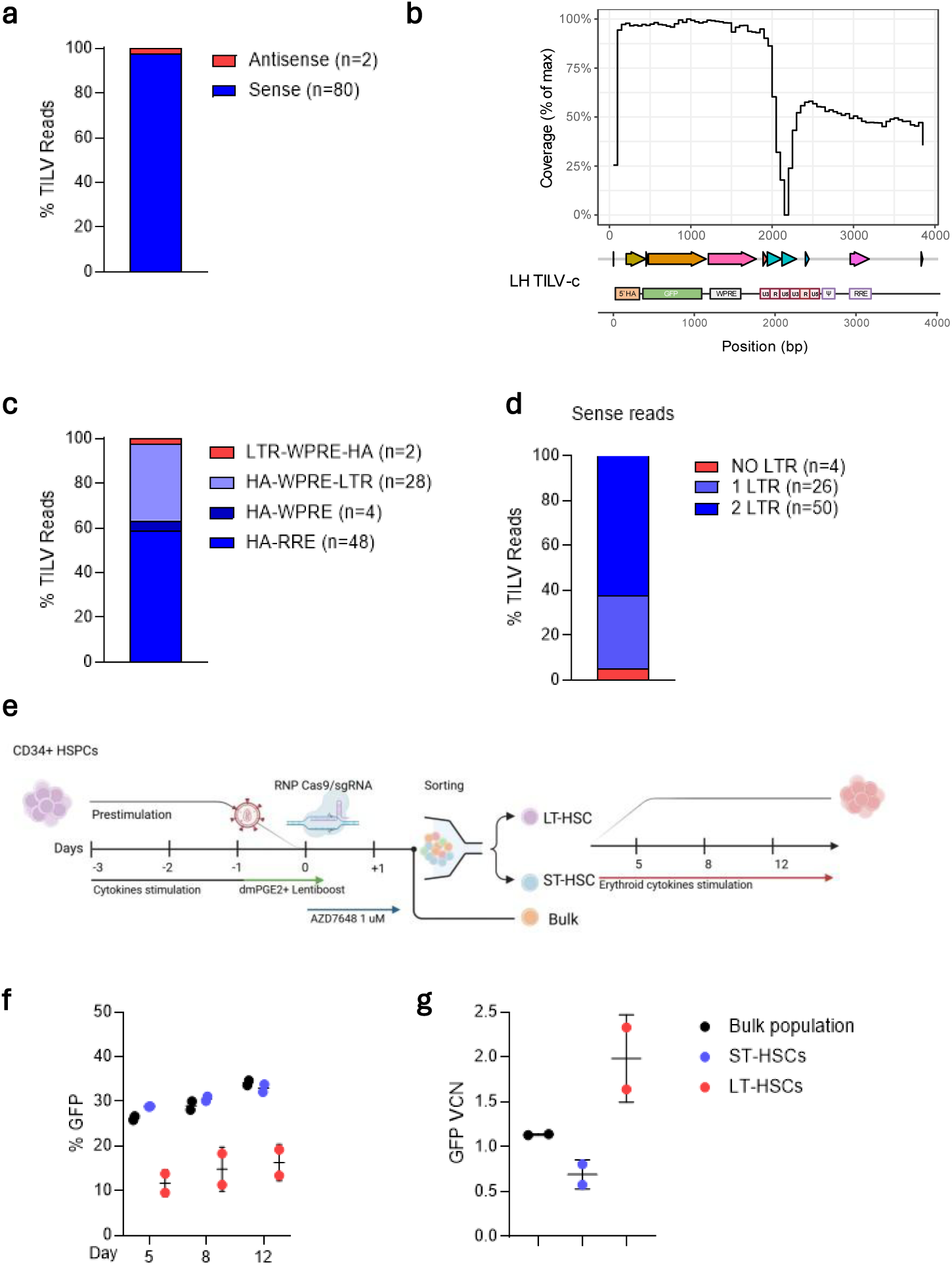
On-target profiling of HSPCs edited with LH-TILV and DNA-PK inhibition. **a-d**) tLRS analysis of LH-TILV edited HSPCs ± 1 µM AZD7648 after 12 days of erythroid differentiation. (a) Orientation of cassette integration (n=86 reads). (b) Read coverage across 4 kb region of chromosome 11 spanning the HBA locus. Below are indicated the different TILV-c elements: LH (long homology), GFP (green fluorescent protein), HIV-1-5’ LRT/HIV-1-3’ LRT (long terminal repeat), Ψ (Psi), WPRE (Woodchuck Hepatitis Virus (WHV) Posttranscriptional Regulatory Element. (c) Analysis of 3’ vector-genome junctions. Integration orientations were assigned as follows: LTR-WPRE-HA (from LTR to HA, antisense), HA-WPRE-LTR (form HA to LTR, without RRE), HA-WPRE (form HA to WPRE, without RRE and LTR), and HA–RRE (from HA to RRE). (d) Number of LTR sequences detected within each tLRS read. **e**) Experimental workflow of HSPC editing and sorting. HSPCs were treated with LH-TILV+AZD7648 1 µM. One day after editing, LT-HSCs (CD34+CD38-CD90+CD45Ra-) and ST-HSCs (CD34+CD38-CD90-CD45Ra-) were sorted and differentiated toward erythroid lineage, together with CD34+ bulk populations. Created with BioRender.com. **f**) Percentage of GFP-positive cells in bulk population, ST- and LT-HSCs at day 5, 8 and 12 of erythroid differentiation (n=2, mean ± SD). **g**) VCN of GFP samples in (f) at day 12 of erythroid differentiation (n=2, mean ± SD).

Finally, we evaluated the feasibility of TILV TI phenotypically defined long-term (LT-) HSCs. Mobilized CD34⁺ cells were edited and subsequently sorted into LT- and short-term (ST-) HSCs (**Fig. 9e; Supplementary Fig. S9c-d).** GFP expression was detected across all subpopulations, though lower fluorescence intensity was observed in LT-HSCs (**Fig. 9f, Supplementary Fig. S9e**), likely reflecting their reduced erythroid differentiation capacity, as indicated by lower CD235 expression (**Supplementary Fig. S9f**). GFP VCN was higher in LT-HSCs (**Fig. 9g**), despite the lower protein expression levels, consistent with enhanced episomal stability in slowly proliferating cells (**Supplementary Fig. S9g**). Additional experiments will determine whether IDLV transduction and/or the integration efficiency are responsible for reduced transgene expression in LT-HSCs.

Altogether, these results demonstrate that combining long homology arm with transient DNA-PK inhibition enables efficient, precise and directional TI of a DNA payload in primary human HSPCs, including long term progenitors, highlighting its potential for future *ex vivo* gene therapy applications.

## Discussion

In this study, we describe TILV (Targeted Integration of Lentiviral Vectors), a CRISPR-assisted strategy that enables precise, directional and efficient targeted gene integration in human HSPCs using integrase-deficient lentiviral vectors (IDLVs) as donor templates.

By embedding an sgRNA target site within TILV cassette, we leverage Cas9 endonuclease activity to linearize episomal IDLV DNA, expose a single homology arm and selectively engage MMEJ/HMEJ repair pathways, resulting in largely seamless, monomeric and orientation-controlled genomic insertions.

We validated the versatility of this approach at the *AAVS1* safe-harbor locus, achieving stable transgene expression under an exogenous promoter, and at the *HBA* locus, demonstrating physiological regulation under an endogenous promoter. Importantly, TILV supports the integration of large DNA cargos (>4 kb, up to ∼9 kb). Compared with conventional lentiviral gene addition, TILV mitigates insertional mutagenesis risk and provides tighter control of the integration site and copy number, allowing predictable expression under exogenous or endogenous promoters. A similar observation was previously reported in T cells; however, no detailed characterization of the donor DNA integration or mechanistic hindsight was provided (47).

A central advantage of TILV lies in the use of IDLV donors, which combine efficient transduction of HSPCs with a packaging capacity (∼10 kb) that overcomes the intrinsic size limit of AAV6 donors (∼4.7 kb). Moreover, IDLV exposure is associated with reduced p53-mediated DNA damage responses compared with AAV6, a factor known to negatively affect HSPC fitness and long-term engraftment potential (27,80).

Vector–genome junction analyses indicate that TILV predominantly relies on homology-mediated end-joining mechanism, resulting in highly precise integrations with minimal indel formation. This efficiency likely arises from the combined use of a single homology arm and template linearization, which favor engagement of the HMEJ pathway (36,81,82).

Accordingly, the improved performance observed with longer homology arm donors is consistent with HMEJ-like processes that benefit from greater resection and homology exposure. Although the mechanistic boundaries between MMEJ, HMEJ and single-strand annealing remain incompletely defined (46,83–85), the junction signatures observed here are consistent with homology-mediated repair operating outside strict S/G2 cell-cycle restriction. This property is particularly relevant for quiescent or slowly cycling HSPCs, which are poorly permissive to HDR-based editing (86). In contrast to NHEJ-based strategies such as HITI, which frequently generate junctional indels (39,41,46,82), TILV achieves high-fidelity, directional TI, although using a single homology arm, in accordance with previous data (81).

Homology arm length and cell type specific repair dynamics likely influence pathway choice While some studies report comparable HMEJ efficiencies across a wide range of homology arm lengths (48 to 500 bp) in activated T cells (36), other studies have shown significant improvements when extending arms from 50 to ∼300 bp in iPSCs (46,79,87). These discrepancies likely reflect intrinsic differences in proliferation and DNA repair profiles among cell types. Our data support the notion that HSPCs benefit from longer homology arms to effectively engage HMEJ, in contrast to highly proliferative cell types, e.g. activated T cells, where these pathways may already be readily accessible (36,46,87).

Targeted long-read sequencing confirmed a predominance of directional orientated, monomeric, full-length cassette insertions, in contrast with the concatemeric or ITR-associated products frequently described for AAV donors (22,88,89). Although occasional integrations of residual lentiviral backbone sequences were detected, the risk associated with these is minimal, as indicated by extensive clinical experience with self-inactivating LV architectures (8,90,91), and can be further reduced by flanking the cassette with two sgRNA target sites to minimize unintended backbone integration.

Pharmacological modulation of DNA repair pathways further enhanced the performance of TILV. Transient DNA-PK inhibition shifted repair away from canonical NHEJ, increasing integration precision and transgene expression in HSPCs, with efficiencies exceeding 40% while preserving multilineage potential. Long-read sequencing confirmed higher utilization CRISPR-based linearized donors and fewer backbone co-integration events, consistent with observations from AAV studies (92,93). Although DNA-PK inhibitors have been associated with increased off-target indels (93), the presence of exogenous donor DNA has been reported to mitigate this effect (74,94). In addition, recent data indicate that DNA-PK inhibition can improve *in vivo* engraftment of gene-edited HSCs, underscoring the potential of combining donor design and pathway modulation to maximize targeting efficiency while maintaining HSPC integrity (95,96).

Other modulators, such as ATR or Polθ inhibitors, merit systematic evaluation to further bias repair toward high-fidelity outcomes while minimizing genomic instability associated with extensive NHEJ disruption (75,76). Additional DNA damage response modulators (e.g., GSE56, Ad5-E4orf6/7, 53BP1 inhibitors) have been shown to enhance AAV TI, editing in LT-HSCs and improve engraftment of edited HSPCs (27,97–100). The systematic evaluation of these synergistic pharmacological interventions for TILV TI will be essential to maximize integration efficiencies while maintaining the highest safety standards for clinical translation.

Despite these advances, two aspects require deeper characterization before clinical translation, the potential for Cas9-independent trapping of donor DNA at endogenous DNA lesions and any residual toxicity associated with IDLV linearization. Additional genome-wide integrations site analysis (101,102) will be necessary to quantify Cas9 independent genomic integration. Likewise, although colony-forming assays and multilineage output suggest a reduced DDR burden relative to AAV-based editing (27,80), direct profiling of p53 activation and damage-response pathways will help validate these observations.

Importantly, TILV system enables therapeutic gene addition under endogenous HBA promoter, as shown for both FVIII and LIPA, resulting in erythroid-specific enzyme secretion and activity. This strategy offers a promising therapeutic avenue for Hemophilia A and Wolman disease, for which curative treatments are still under development (50), and in general, for protein replacement and cross-correction strategies in lysosomal storage disorders.

Finally, achieving TI in LT-HSCs remains a major bottleneck for genome editing. Our preliminary results indicate that TILV can reach primitive HSCs at levels comparable to AAV6-HDR (27), warranting future long-term xenograft studies to assess durability and repopulation capacity.

Additional improvements in HSPCs transduction, through optimized LV pseudotypes, envelope engineering (103–105) or non-electroporation-based RNP delivery systems (lipid nanoparticles (36); virus-like particles (106,107); peptide mediated delivery (108–110) may further increase access to quiescent HSCs and enable future *in vivo* applications.

Alternative non-viral donors, such as linear and circular ssDNA and nanoplasmids, have demonstrated the ability to support efficient editing and stable integration of large cargos in T cells (36,111–115), but their applicability in HSPCs remains limited. Similarly, emerging DSB-free gene insertion systems (116–121) aim to achieve TI independently of the cell cycle; yet, most exhibit low efficiency and have not been evaluated in HSPCs. For now, HDR-based approaches remain the most established strategies for precise genome editing in human HSPCs.

In summary, TILV provides an efficient and accurate platform for targeted gene addition in human HSPCs by combining CRISPR-directed donor linearization with (micro)homology-mediated repair pathways. This approach addresses key limitations of HDR-based editing while maintaining large cargo capability and reduced cytotoxicity, establishing TILV as a versatile and promising platform for next-generation gene and cell therapies, with broad applicability for durable protein replacement and cross-correction strategies.

### Statistical analysis

Statistical analyses were performed using GraphPad Prism version 10.00 for Windows (GraphPad Software, La Jolla, CA, USA, “www.graphpad.com<u>”</u>). Chi-square test, Exact Binomial Test, Paired or Unpaired t-test, two-way ANOVA were performed as indicated. Values are expressed as mean ± standard deviation (SD). Percentage data were analyzed using variance-stabilizing transformations when required.

## Data availability

The authors declare that all data supporting the findings of this study are available in the main text or the supplementary materials. The Guide-seq, Amplicon-seq and LRS data that support the findings of this study have been deposited (https://dataview.ncbi.nlm.nih.gov/object/PRJNA1442083?reviewer=2pk7j8t1fk03s2qpv2hikjol0 <u>b</u>). Any other relevant data is available at reasonable request.

## Acknowledgments

We acknowledge the Genethon “Vector Core Facility” for their help with IDLV production, Genethon Platform “Imaging and Cytometry Core Facility” for cell sorting and FACS technical assistance. We acknowledge Dr J.P. Concordet for Cas9 protein production (supported by ANR-II-INSB-0014 and France 2030 PEPR BBIT Edito 22-PEBI-0005, INSERM U1154). We thank members of the “Therapeutic genome editing” group for fruitful discussions and experimental help. We are grateful to Ile-de-France Region, to Conseil Departemental de l’Essonne (ASTRE), INSERM and GIP Genopole Evry for the purchase of the equipment.

## Authors’ contributions

Giulia Scalisi: Conceived the study, designed and performed experiments, analyzed data and wrote the original draft of the manuscript. Aboud Sakkal: Designed and performed experiments and analyzed data. Laurie Lacombe, Francesco Sarnari and Michela Rosiello: Performed experiments and analyzed data. Paola Galbiati and Marine Rouillon: Performed Amplicon sequencing and CRISPResso analysis. Marie As: produced Cas9 protein. Marcello Maresca: Conceived and Mike Firth: Performed RIMA analysis. Guillaume Core and Alexandra Tachtsidi: Performed targeted long read sequencing and bioinformatics analysis. Ivan Peyron and Peter J. Leting: Contributed to FVIII experimental analysis. Julie Oustelandt and Marine Laurent contributed to LAL experiments. Giulia Pavani contributed to initial plasmid design and supervised selected experiments. Anne Galy and Annarita Miccio: Provided input on study design and data analysis. Mario Amendola: Conceived and supervised the study and wrote the manuscript.

## Funding

This work was funded by the European Union (EDITSCD grant 101057659, https://editscd.eu/ and MAGIC grant 101080690, https://magic-horizon.eu; ERDERA grant 101156595, https://erdera.org/). M.A. was also funded by the Genethon, the AFM-Telethon foundation (BE-DREP grant), INSERM, the University of Evry Val d’Essonne, the French National Research Agency (grants: NEEDED ANR-24-CE18-3712-01; PEMGeT ANR-22-CE17-0028-02; HemoLen ANR-20-CE17-0016-01; IRIS ANR-21-CE14-0063-03), the France Relance program, the DIM Therapie Genique (SafeSCD), The views and opinions expressed herein are those of the authors only and do not necessarily reflect those of the European Union or HaDEA. Neither the European Union nor the granting authorities can be held responsible for them.

## Conflict of interest

M.A, A.S and G.P. are inventors on a patent application related to this work filed by Genethon (PCT/EP2020/070622, filed 22 July 2022). All other authors declare that they have no competing or financial interests.

## Declaration of generative AI and AI-assisted technologies in the writing process

During the preparation of this work, the authors used generative AI tools to improve clarity and streamline the text to meet the journal’s space constraints. All scientific content, data interpretation and conclusions were critically reviewed and edited by the authors, who take full responsibility for the integrity and accuracy of the published work.

## Notes

### Competing Interest Statement

The authors have declared no competing interest.

## Reference

1. Doudna JA. The promise and challenge of therapeutic genome editing. Nature. 2020 Feb 13;578(7794):229–36. doi:10.1038/s41586-020-1978-5

2. Hubbard N, Hagin D, Sommer K, Song Y, Khan I, Clough C, et al. Targeted gene editing restores regulated CD40L function in X-linked hyper-IgM syndrome. Blood. 2016 May 26;127(21):2513–22. doi:10.1182/blood-2015-11-683235

3. Ferrari S, Valeri E, Conti A, Scala S, Aprile A, Di Micco R, et al. Genetic engineering meets hematopoietic stem cell biology for next-generation gene therapy. Cell Stem Cell. 2023 May;30(5):549–70. doi:10.1016/j.stem.2023.04.014

4. Kohn DB, Booth C, Shaw KL, Xu-Bayford J, Garabedian E, Trevisan V, et al. Autologous Ex Vivo Lentiviral Gene Therapy for Adenosine Deaminase Deficiency. N Engl J Med. 2021 May 27;384(21):2002–13. doi:10.1056/NEJMoa2027675 PubMed PMID: 33974366; PubMed Central PMCID: PMC8240285.

5. Magnani A, Semeraro M, Adam F, Booth C, Dupré L, Morris EC, et al. Author Correction: Long-term safety and efficacy of lentiviral hematopoietic stem/progenitor cell gene therapy for Wiskott-Aldrich syndrome. Nat Med. 2022 Oct;28(10):2217. doi:10.1038/s41591-022-01985-y PubMed PMID: 35945284; PubMed Central PMCID: PMC9556294.

7. Schröder ARW, Shinn P, Chen H, Berry C, Ecker JR, Bushman F. HIV-1 integration in the human genome favors active genes and local hotspots. Cell. 2002 Aug 23;110(4):521–9. doi:10.1016/s0092-8674(02)00864-4 PubMed PMID: 12202041.

8. Montini E, Naldini L, Booth C, Kohn DB, Aiuti A. Balancing efficacy and safety in lentiviral vector-mediated hematopoietic stem cell gene therapy. Mol Ther. 2025 Jan 8;33(1):6–8. doi:10.1016/j.ymthe.2024.12.028 PubMed PMID: 39729983; PubMed Central PMCID: PMC11764623.

9. Tucci F, Galimberti S, Naldini L, Valsecchi MG, Aiuti A. A systematic review and meta-analysis of gene therapy with hematopoietic stem and progenitor cells for monogenic disorders. Nat Commun. 2022 Mar 14;13(1):1315. doi:10.1038/s41467-022-28762-2 PubMed PMID: 35288539; PubMed Central PMCID: PMC8921234.

10. Duncan CN, Bledsoe JR, Grzywacz B, Beckman A, Bonner M, Eichler FS, et al. Hematologic Cancer after Gene Therapy for Cerebral Adrenoleukodystrophy. N Engl J Med. 2024 Oct 10;391(14):1287–301. doi:10.1056/NEJMoa2405541 PubMed PMID: 39383458; PubMed Central PMCID: PMC11846662.

11. Chen HC, Martinez JP, Zorita E, Meyerhans A, Filion GJ. Position effects influence HIV latency reversal. Nat Struct Mol Biol. 2017 Jan;24(1):47–54. doi:10.1038/nsmb.3328 PubMed PMID: 27870832.

12. Vansant G, Chen HC, Zorita E, Trejbalová K, Miklík D, Filion G, et al. The chromatin landscape at the HIV-1 provirus integration site determines viral expression. Nucleic Acids Research. 2020 Aug 20;48(14):7801–17. doi:10.1093/nar/gkaa536

13. Akhtar W, de Jong J, Pindyurin AV, Pagie L, Meuleman W, de Ridder J, et al. Chromatin position effects assayed by thousands of reporters integrated in parallel. Cell. 2013 Aug 15;154(4):914–27. doi:10.1016/j.cell.2013.07.018 PubMed PMID: 23953119.

14. Maricque BB, Chaudhari HG, Cohen BA. A massively parallel reporter assay dissects the influence of chromatin structure on cis-regulatory activity. Nat Biotechnol. 2018 Nov 19. doi:10.1038/nbt.4285 PubMed PMID: 30451991; PubMed Central PMCID: PMC7351048.

15. Ferris AL, Wu X, Hughes CM, Stewart C, Smith SJ, Milne TA, et al. Lens epithelium-derived growth factor fusion proteins redirect HIV-1 DNA integration. Proc Natl Acad Sci U S A. 2010 Feb 16;107(7):3135–40. doi:10.1073/pnas.0914142107 PubMed PMID: 20133638; PubMed Central PMCID: PMC2840313.

16. Gijsbers R, Ronen K, Vets S, Malani N, De Rijck J, McNeely M, et al. LEDGF hybrids efficiently retarget lentiviral integration into heterochromatin. Mol Ther. 2010 Mar;18(3):552–60. doi:10.1038/mt.2010.36 PubMed PMID: 20195265; PubMed Central PMCID: PMC2839429.

17. Vink CA, Gaspar HB, Gabriel R, Schmidt M, McIvor RS, Thrasher AJ, et al. Sleeping beauty transposition from nonintegrating lentivirus. Mol Ther. 2009 Jul;17(7):1197–204. doi:10.1038/mt.2009.94 PubMed PMID: 19417741; PubMed Central PMCID: PMC2835211.

18. Thomsen EA, Skipper KA, Andersen S, Haslund D, Skov TW, Mikkelsen JG. CRISPR-Cas9-directed gene tagging using a single integrase-defective lentiviral vector carrying a transposase-based Cas9 off switch. Mol Ther Nucleic Acids. 2022 Sep 13;29:563–76. doi:10.1016/j.omtn.2022.08.005 PubMed PMID: 36090759; PubMed Central PMCID: PMC9403905.

19. Lombardo A, Cesana D, Genovese P, Di Stefano B, Provasi E, Colombo DF, et al. Site-specific integration and tailoring of cassette design for sustainable gene transfer. Nat Methods. 2011 Oct;8(10):861–9. doi:10.1038/nmeth.1674

20. Voit RA, Hendel A, Pruett-Miller SM, Porteus MH. Nuclease-mediated gene editing by homologous recombination of the human globin locus. Nucleic Acids Research. 2014 Jan 1;42(2):1365–78. doi:10.1093/nar/gkt947

21. Hubbard BP, Badran AH, Zuris JA, Guilinger JP, Davis KM, Chen L, et al. Continuous directed evolution of DNA-binding proteins to improve TALEN specificity. Nat Methods. 2015 Oct;12(10):939–42. doi:10.1038/nmeth.3515 PubMed PMID: 26258293; PubMed Central PMCID: PMC4589463.

22. Schiroli G, Ferrari S, Conway A, Jacob A, Capo V, Albano L, et al. Preclinical modeling highlights the therapeutic potential of hematopoietic stem cell gene editing for correction of SCID-X1. Sci Transl Med. 2017 Oct 11;9(411):eaan0820. doi:10.1126/scitranslmed.aan0820

23. Dever DP, Bak RO, Reinisch A, Camarena J, Washington G, Nicolas CE, et al. CRISPR/Cas9 β-globin gene targeting in human haematopoietic stem cells. Nature. 2016 Nov 17;539(7629):384–9. doi:10.1038/nature20134

24. Wang J, Exline CM, DeClercq JJ, Llewellyn GN, Hayward SB, Li PWL, et al. Homology-driven genome editing in hematopoietic stem and progenitor cells using ZFN mRNA and AAV6 donors. Nat Biotechnol. 2015 Dec;33(12):1256–63. doi:10.1038/nbt.3408

25. Pavani G, Laurent M, Fabiano A, Cantelli E, Sakkal A, Corre G, et al. Ex vivo editing of human hematopoietic stem cells for erythroid expression of therapeutic proteins. Nat Commun. 2020 Jul 29;11(1):3778. doi:10.1038/s41467-020-17552-3

26. Pavani G, Fabiano A, Laurent M, Amor F, Cantelli E, Chalumeau A, et al. Correction of β-thalassemia by CRISPR/Cas9 editing of the α-globin locus in human hematopoietic stem cells. Blood Advances. 2021 Mar 9;5(5):1137–53. doi:10.1182/bloodadvances.2020001996

27. Ferrari S, Jacob A, Cesana D, Laugel M, Beretta S, Varesi A, et al. Choice of template delivery mitigates the genotoxic risk and adverse impact of editing in human hematopoietic stem cells. Cell Stem Cell. 2022 Oct;29(10):1428–1444.e9. doi:10.1016/j.stem.2022.09.001

28. Conti A, Giannetti K, Midena F, Beretta S, Gualandi N, De Marco R, et al. Senescence and inflammation are unintended adverse consequences of CRISPR-Cas9/AAV6-mediated gene editing in hematopoietic stem cells. Cell Reports Medicine. 2025 Jun;6(6):102157. doi:10.1016/j.xcrm.2025.102157

29. San Filippo J, Sung P, Klein H. Mechanism of eukaryotic homologous recombination. Annu Rev Biochem. 2008;77:229–57. doi:10.1146/annurev.biochem.77.061306.125255 PubMed PMID: 18275380.

30. Gaj T, Staahl BT, Rodrigues GMC, Limsirichai P, Ekman FK, Doudna JA, et al. Targeted gene knock-in by homology-directed genome editing using Cas9 ribonucleoprotein and AAV donor delivery. Nucleic Acids Research. 2017 Jun 20;45(11):e98–e98. doi:10.1093/nar/gkx154

31. Baker O, Tsurkan S, Fu J, Klink B, Rump A, Obst M, et al. The contribution of homology arms to nuclease-assisted genome engineering. Nucleic Acids Research. 2017 Jul 27;45(13):8105–15. doi:10.1093/nar/gkx497

32. Ling C, Yu C, Wang C, Yang M, Yang H, Yang K, et al. rAAV capsid mutants eliminate leaky expression from DNA donor template for homologous recombination. Nucleic Acids Res. 2024 Jun 24;52(11):6518–31. doi:10.1093/nar/gkae401 PubMed PMID: 38783157; PubMed Central PMCID: PMC11194064.

33. Lalanne JB, Mich JK, Huynh C, Hunker AC, McDiarmid TA, Levi BP, et al. Extensive length and homology dependent chimerism in pool-packaged AAV libraries [Internet]. Genomics; 2025 [cited 2026 May 2]. Available from: http://biorxiv.org/lookup/doi/10.1101/2025.01.14.632594 doi:10.1101/2025.01.14.632594

34. Taran JA, Mintaev RR, Glazkova DV, Belugin BV, Bogoslovskaya EV, Shipulin GA. [Influence of Homology Arm Length and Structure on the Efficiency of Long Transgene Integration into a Cleavage Site Induced by SpCas9 or AsCpf1]. Mol Biol (Mosk). 2025;59(2):255–65. doi:10.31857/S0026898425020079 PubMed PMID: 40558037.

35. Deyle DR, Li LB, Ren G, Russell DW. The effects of polymorphisms on human gene targeting. Nucleic Acids Res. 2014 Mar;42(5):3119–24. doi:10.1093/nar/gkt1303 PubMed PMID: 24371280; PubMed Central PMCID: PMC3950700.

36. Webber BR, Johnson MJ, Skeate JG, Slipek NJ, Lahr WS, DeFeo AP, et al. Cas9-induced targeted integration of large DNA payloads in primary human T cells via homology-mediated end-joining DNA repair. Nat Biomed Eng. 2023 Dec 13. doi:10.1038/s41551-023-01157-4

37. Maresca M, Lin VG, Guo N, Yang Y. Obligate ligation-gated recombination (ObLiGaRe): custom-designed nuclease-mediated targeted integration through nonhomologous end joining. Genome Res. 2013 Mar;23(3):539–46. doi:10.1101/gr.145441.112 PubMed PMID: 23152450; PubMed Central PMCID: PMC3589542.

38. Suzuki K, Tsunekawa Y, Hernandez-Benitez R, Wu J, Zhu J, Kim EJ, et al. In vivo genome editing via CRISPR/Cas9 mediated homology-independent targeted integration. Nature. 2016 Dec 1;540(7631):144–9. doi:10.1038/nature20565

39. Bloomer H, Smith RH, Hakami W, Larochelle A. Genome editing in human hematopoietic stem and progenitor cells via CRISPR-Cas9-mediated homology-independent targeted integration. Molecular Therapy. 2021 Apr;29(4):1611–24. doi:10.1016/j.ymthe.2020.12.010

40. Tornabene P, Ferla R, Llado-Santaeularia M, Centrulo M, Dell’Anno M, Esposito F, et al. Therapeutic homology-independent targeted integration in retina and liver. Nat Commun. 2022 Apr 12;13(1):1963. doi:10.1038/s41467-022-29550-8

41. Esposito F, Dell’Aquila F, Rhiel M, Auricchio S, Chmielewski KO, Andrieux G, et al. Safe and effective liver-directed AAV-mediated homology-independent targeted integration in mouse models of inherited diseases. Cell Reports Medicine. 2024 Jul;5(7):101619. doi:10.1016/j.xcrm.2024.101619

42. Nakade S, Tsubota T, Sakane Y, Kume S, Sakamoto N, Obara M, et al. Microhomology-mediated end-joining-dependent integration of donor DNA in cells and animals using TALENs and CRISPR/Cas9. Nat Commun. 2014 Nov 20;5(1):5560. doi:10.1038/ncomms6560

43. Sakuma T, Nakade S, Sakane Y, Suzuki KIT, Yamamoto T. MMEJ-assisted gene knock-in using TALENs and CRISPR-Cas9 with the PITCh systems. Nat Protoc. 2016 Jan;11(1):118–33. doi:10.1038/nprot.2015.140

44. Decottignies A. Microhomology-mediated end joining in fission yeast is repressed by pku70 and relies on genes involved in homologous recombination. Genetics. 2007 Jul;176(3):1403–15. doi:10.1534/genetics.107.071621 PubMed PMID: 17483423; PubMed Central PMCID: PMC1931558.

45. Truong LN, Li Y, Shi LZ, Hwang PYH, He J, Wang H, et al. Microhomology-mediated End Joining and Homologous Recombination share the initial end resection step to repair DNA double-strand breaks in mammalian cells. Proc Natl Acad Sci U S A. 2013 May 7;110(19):7720–5. doi:10.1073/pnas.1213431110 PubMed PMID: 23610439; PubMed Central PMCID: PMC3651503.

46. Yao X, Wang X, Hu X, Liu Z, Liu J, Zhou H, et al. Homology-mediated end joining-based targeted integration using CRISPR/Cas9. Cell Res. 2017 Jun;27(6):801–14. doi:10.1038/cr.2017.76

47. Chavez M, Rane DA, Chen X, Qi LS. Stable expression of large transgenes via the knock-in of an integrase-deficient lentivirus. Nat Biomed Eng. 2023 May 1;7(5):661–71. doi:10.1038/s41551-023-01037-x

48. Gilpatrick T, Lee I, Graham JE, Raimondeau E, Bowen R, Heron A, et al. Targeted nanopore sequencing with Cas9-guided adapter ligation. Nat Biotechnol. 2020 Apr;38(4):433–8. doi:10.1038/s41587-020-0407-5 PubMed PMID: 32042167; PubMed Central PMCID: PMC7145730.

49. Montague TG, Cruz JM, Gagnon JA, Church GM, Valen E. CHOPCHOP: a CRISPR/Cas9 and TALEN web tool for genome editing. Nucleic Acids Res. 2014 Jul;42(Web Server issue):W401–407. doi:10.1093/nar/gku410 PubMed PMID: 24861617; PubMed Central PMCID: PMC4086086.

50. Laurent M, Harb R, Jenny C, Oustelandt J, Jimenez S, Cosette J, et al. Rescue of lysosomal acid lipase deficiency in mice by rAAV8 liver gene transfer. Commun Med (Lond). 2025 Apr 11;5(1):110. doi:10.1038/s43856-025-00816-8 PubMed PMID: 40216942; PubMed Central PMCID: PMC11992068.

51. Laurent M, Cosette J, Pavani G, Bayol S, Jenny C, Harb R, et al. Advanced Imaging and Cytometric Techniques to Characterize Lipid Accumulation in Wolman Disease. Cytometry A. 2025 Jul;107(7):464–75. doi:10.1002/cyto.a.24949 PubMed PMID: 40613725.

52. McIntosh J, Lenting PJ, Rosales C, Lee D, Rabbanian S, Raj D, et al. Therapeutic levels of FVIII following a single peripheral vein administration of rAAV vector encoding a novel human factor VIII variant. Blood. 2013 Apr 25;121(17):3335–44. doi:10.1182/blood-2012-10-462200

53. Siner JI, Iacobelli NP, Sabatino DE, Ivanciu L, Zhou S, Poncz M, et al. Minimal modification in the factor VIII B-domain sequence ameliorates the murine hemophilia A phenotype. Blood. 2013 May 23;121(21):4396–403. doi:10.1182/blood-2012-10-464164 PubMed PMID: 23372167; PubMed Central PMCID: PMC3663432.

54. Amendola M, Venneri MA, Biffi A, Vigna E, Naldini L. Coordinate dual-gene transgenesis by lentiviral vectors carrying synthetic bidirectional promoters. Nat Biotechnol. 2005 Jan;23(1):108–16. doi:10.1038/nbt1049 PubMed PMID: 15619618.

55. Conant D, Hsiau T, Rossi N, Oki J, Maures T, Waite K, et al. Inference of CRISPR Edits from Sanger Trace Data. The CRISPR Journal. 2022 Feb 1;5(1):123–30. doi:10.1089/crispr.2021.0113

56. Brinkman EK, Chen T, Amendola M, van Steensel B. Easy quantitative assessment of genome editing by sequence trace decomposition. Nucleic Acids Research. 2014 Dec 16;42(22):e168–e168. doi:10.1093/nar/gku936

57. Clement K, Rees H, Canver MC, Gehrke JM, Farouni R, Hsu JY, et al. CRISPResso2 provides accurate and rapid genome editing sequence analysis. Nat Biotechnol. 2019 Mar;37(3):224–6. doi:10.1038/s41587-019-0032-3

58. Taheri-Ghahfarokhi A, Taylor BJM, Nitsch R, Lundin A, Cavallo AL, Madeyski-Bengtson K, et al. Decoding non-random mutational signatures at Cas9 targeted sites. Nucleic Acids Research. 2018 Sep 19;46(16):8417–34. doi:10.1093/nar/gky653

59. Tsai SQ, Zheng Z, Nguyen NT, Liebers M, Topkar VV, Thapar V, et al. GUIDE-seq enables genome-wide profiling of off-target cleavage by CRISPR-Cas nucleases. Nat Biotechnol. 2015 Feb;33(2):187–97. doi:10.1038/nbt.3117 PubMed PMID: 25513782; PubMed Central PMCID: PMC4320685.

60. Corre G, Rouillon M, Mombled M, Amendola M. Advanced pipeline for CRISPR/Cas9 off-targets detection in Guide-seq and related integration-based assays [Internet]. Bioinformatics; 2025 [cited 2026 May 2]. Available from: http://biorxiv.org/lookup/doi/10.1101/2025.09.30.679427 doi:10.1101/2025.09.30.679427

61. Munir S, Thierry S, Subra F, Deprez E, Delelis O. Quantitative analysis of the time-course of viral DNA forms during the HIV-1 life cycle. Retrovirology. 2013 Dec;10(1):87. doi:10.1186/1742-4690-10-87

62. Wanisch K, Yáñez-Muñoz RJ. Integration-deficient Lentiviral Vectors: A Slow Coming of Age. Molecular Therapy. 2009 Aug;17(8):1316–32. doi:10.1038/mt.2009.122

63. Concordet JP, Haeussler M. CRISPOR: intuitive guide selection for CRISPR/Cas9 genome editing experiments and screens. Nucleic Acids Research. 2018 Jul 2;46(W1):W242–5. doi:10.1093/nar/gky354

64. Cradick TJ, Qiu P, Lee CM, Fine EJ, Bao G. COSMID: A Web-based Tool for Identifying and Validating CRISPR/Cas Off-target Sites. Mol Ther Nucleic Acids. 2014 Dec 2;3(12):e214. doi:10.1038/mtna.2014.64 PubMed PMID: 25462530; PubMed Central PMCID: PMC4272406.

65. Cromer MK, Camarena J, Martin RM, Lesch BJ, Vakulskas CA, Bode NM, et al. Gene replacement of α-globin with β-globin restores hemoglobin balance in β-thalassemia-derived hematopoietic stem and progenitor cells. Nat Med. 2021 Apr;27(4):677–87. doi:10.1038/s41591-021-01284-y PubMed PMID: 33737751; PubMed Central PMCID: PMC8265212.

66. Heffner GC, Bonner M, Christiansen L, Pierciey FJ, Campbell D, Smurnyy Y, et al. Prostaglandin E2 Increases Lentiviral Vector Transduction Efficiency of Adult Human Hematopoietic Stem and Progenitor Cells. Molecular Therapy. 2018 Jan;26(1):320–8. doi:10.1016/j.ymthe.2017.09.025

67. Poletti V, Montepeloso A, Pellin D, Biffi A. Prostaglandin E2 as transduction enhancer affects competitive engraftment of human hematopoietic stem and progenitor cells. Molecular Therapy Methods & Clinical Development. 2023 Dec;31:101131. doi:10.1016/j.omtm.2023.101131

68. Hauber I, Beschorner N, Schrödel S, Chemnitz J, Kröger N, Hauber J, et al. Improving Lentiviral Transduction of CD34^+^ Hematopoietic Stem and Progenitor Cells. Human Gene Therapy Methods. 2018 Apr;29(2):104–13. doi:10.1089/hgtb.2017.085

69. Jang Y, Kim YS, Wielgosz MM, Ferrara F, Ma Z, Condori J, et al. Optimizing lentiviral vector transduction of hematopoietic stem cells for gene therapy. Gene Ther. 2020 Dec;27(12):545–56. doi:10.1038/s41434-020-0150-z

70. Petrillo C, Thorne LG, Unali G, Schiroli G, Giordano AMS, Piras F, et al. Cyclosporine H Overcomes Innate Immune Restrictions to Improve Lentiviral Transduction and Gene Editing In Human Hematopoietic Stem Cells. Cell Stem Cell. 2018 Dec;23(6):820–832.e9. doi:10.1016/j.stem.2018.10.008

71. Wang CX, Sather BD, Wang X, Adair J, Khan I, Singh S, et al. Rapamycin relieves lentiviral vector transduction resistance in human and mouse hematopoietic stem cells. Blood. 2014 Aug 7;124(6):913–23. doi:10.1182/blood-2013-12-546218 PubMed PMID: 24914132; PubMed Central PMCID: PMC4126331.

72. Robert F, Barbeau M, Éthier S, Dostie J, Pelletier J. Pharmacological inhibition of DNA-PK stimulates Cas9-mediated genome editing. Genome Med. 2015 Dec;7(1):93. doi:10.1186/s13073-015-0215-6

73. Schimmel J, Muñoz-Subirana N, Kool H, van Schendel R, van der Vlies S, Kamp JA, et al. Modulating mutational outcomes and improving precise gene editing at CRISPR-Cas9-induced breaks by chemical inhibition of end-joining pathways. Cell Reports. 2023 Feb;42(2):112019. doi:10.1016/j.celrep.2023.112019

74. Riesenberg S, Chintalapati M, Macak D, Kanis P, Maricic T, Pääbo S. Simultaneous precise editing of multiple genes in human cells. Nucleic Acids Research. 2019 Nov 4;47(19):e116–e116. doi:10.1093/nar/gkz669

75. Selvaraj S, Feist WN, Viel S, Vaidyanathan S, Dudek AM, Gastou M, et al. High-efficiency transgene integration by homology-directed repair in human primary cells using DNA-PKcs inhibition. Nat Biotechnol. 2024 May;42(5):731–44. doi:10.1038/s41587-023-01888-4

76. Wimberger S, Akrap N, Firth M, Brengdahl J, Engberg S, Schwinn MK, et al. Simultaneous inhibition of DNA-PK and Polϴ improves integration efficiency and precision of genome editing. Nat Commun. 2023 Aug 14;14(1):4761. doi:10.1038/s41467-023-40344-4

77. Fok JHL, Ramos-Montoya A, Vazquez-Chantada M, Wijnhoven PWG, Follia V, James N, et al. AZD7648 is a potent and selective DNA-PK inhibitor that enhances radiation, chemotherapy and olaparib activity. Nat Commun. 2019 Nov 7;10(1):5065. doi:10.1038/s41467-019-12836-9

78. Benitez EK, Lomova Kaufman A, Cervantes L, Clark DN, Ayoub PG, Senadheera S, et al. Global and Local Manipulation of DNA Repair Mechanisms to Alter Site-Specific Gene Editing Outcomes in Hematopoietic Stem Cells. Front Genome Ed. 2020 Dec 10;2:601541. doi:10.3389/fgeed.2020.601541

79. Gray DH, Santos J, Keir AG, Villegas I, Maddock S, Trope EC, et al. A comparison of DNA repair pathways to achieve a site-specific gene modification of the Bruton’s tyrosine kinase gene. Molecular Therapy Nucleic Acids. 2022 Mar;27:505–16. doi:10.1016/j.omtn.2021.12.014

80. Schiroli G, Conti A, Ferrari S, Della Volpe L, Jacob A, Albano L, et al. Precise Gene Editing Preserves Hematopoietic Stem Cell Function following Transient p53-Mediated DNA Damage Response. Cell Stem Cell. 2019 Apr 4;24(4):551–565.e8. doi:10.1016/j.stem.2019.02.019 PubMed PMID: 30905619; PubMed Central PMCID: PMC6458988.

81. Suzuki K, Yamamoto M, Hernandez-Benitez R, Li Z, Wei C, Soligalla RD, et al. Precise in vivo genome editing via single homology arm donor mediated intron-targeting gene integration for genetic disease correction. Cell Res. 2019 Oct;29(10):804–19. doi:10.1038/s41422-019-0213-0

82. Jin X, Wu X, Song J, Luo M, Ye Q, Ren C, et al. Comparative evaluation of liver-directed knockin strategies with viral and nonviral vectors in mouse inherited disease models. Mol Ther. 2026 Mar 4;34(3):1775–93. doi:10.1016/j.ymthe.2025.12.012 PubMed PMID: 41376155; PubMed Central PMCID: PMC12974188.

83. Ceccaldi R, Rondinelli B, D’Andrea AD. Repair Pathway Choices and Consequences at the Double-Strand Break. Trends Cell Biol. 2016 Jan;26(1):52–64. doi:10.1016/j.tcb.2015.07.009 PubMed PMID: 26437586; PubMed Central PMCID: PMC4862604.

84. Wierson WA, Welker JM, Almeida MP, Mann CM, Webster DA, Torrie ME, et al. Efficient targeted integration directed by short homology in zebrafish and mammalian cells. Elife. 2020 May 15;9:e53968. doi:10.7554/eLife.53968 PubMed PMID: 32412410; PubMed Central PMCID: PMC7228771.

85. Ruis BL, Bielinsky AK, Hendrickson EA. Gene editing and CRISPR-dependent homology-mediated end joining. Exp Mol Med. 2025 Jul;57(7):1409–18. doi:10.1038/s12276-025-01442-z PubMed PMID: 40745005; PubMed Central PMCID: PMC12322213.

86. Genovese P, Schiroli G, Escobar G, Di Tomaso T, Firrito C, Calabria A, et al. Targeted genome editing in human repopulating haematopoietic stem cells. Nature. 2014 Jun;510(7504):235–40. doi:10.1038/nature13420

87. Zhang JP, Li XL, Li GH, Chen W, Arakaki C, Botimer GD, et al. Efficient precise knockin with a double cut HDR donor after CRISPR/Cas9-mediated double-stranded DNA cleavage. Genome Biol. 2017 Feb 20;18(1):35. doi:10.1186/s13059-017-1164-8

88. Suchy FP, Karigane D, Nakauchi Y, Higuchi M, Zhang J, Pekrun K, et al. Genome engineering with Cas9 and AAV repair templates generates frequent concatemeric insertions of viral vectors. Nat Biotechnol. 2025 Feb;43(2):204–13. doi:10.1038/s41587-024-02171-w PubMed PMID: 38589662; PubMed Central PMCID: PMC11524221.

89. Simpson BP, Yrigollen CM, Izda A, Davidson BL. Targeted long-read sequencing captures CRISPR editing and AAV integration outcomes in brain. Mol Ther. 2023 Mar 1;31(3):760–73. doi:10.1016/j.ymthe.2023.01.004 PubMed PMID: 36617193; PubMed Central PMCID: PMC10014281.

90. Dull T, Zufferey R, Kelly M, Mandel RJ, Nguyen M, Trono D, et al. A third-generation lentivirus vector with a conditional packaging system. J Virol. 1998 Nov;72(11):8463–71. doi:10.1128/JVI.72.11.8463-8471.1998 PubMed PMID: 9765382; PubMed Central PMCID: PMC110254.

91. Cesana D, Ranzani M, Volpin M, Bartholomae C, Duros C, Artus A, et al. Uncovering and dissecting the genotoxicity of self-inactivating lentiviral vectors in vivo. Mol Ther. 2014 Apr;22(4):774–85. doi:10.1038/mt.2014.3 PubMed PMID: 24441399; PubMed Central PMCID: PMC3982501.

92. Dalwadi DA, Calabria A, Tiyaboonchai A, Posey J, Naugler WE, Montini E, et al. AAV integration in human hepatocytes. Mol Ther. 2021 Oct 6;29(10):2898–909. doi:10.1016/j.ymthe.2021.08.031 PubMed PMID: 34461297; PubMed Central PMCID: PMC8531150.

93. Cullot G, Aird EJ, Schlapansky MF, Yeh CD, van de Venn L, Vykhlyantseva I, et al. Genome editing with the HDR-enhancing DNA-PKcs inhibitor AZD7648 causes large-scale genomic alterations. Nat Biotechnol. 2025 Nov;43(11):1778–82. doi:10.1038/s41587-024-02488-6 PubMed PMID: 39604565; PubMed Central PMCID: PMC12611759.

94. Wen W, Quan ZJ, Li SA, Yang ZX, Fu YW, Zhang F, et al. Effective control of large deletions after double-strand breaks by homology-directed repair and dsODN insertion. Genome Biol. 2021 Aug 20;22(1):236. doi:10.1186/s13059-021-02462-4 PubMed PMID: 34416913; PubMed Central PMCID: PMC8377869.

95. Pugliano CM, Berger M, Ray RM, Sapkos K, Wu B, Laird A, et al. DNA-PK inhibition enhances gene editing efficiency in HSPCs for CRISPR-based treatment of X-linked hyper IgM syndrome. Mol Ther Methods Clin Dev. 2024 Sep 12;32(3):101297. doi:10.1016/j.omtm.2024.101297 PubMed PMID: 40012884; PubMed Central PMCID: PMC11863497.

96. Dudek AM, Feist WN, Sasu EJ, Luna SE, Ben-Efraim K, Bak RO, et al. A simultaneous knockout knockin genome editing strategy in HSPCs potently inhibits CCR5- and CXCR4-tropic HIV-1 infection. Cell Stem Cell. 2024 Apr 4;31(4):499–518.e6. doi:10.1016/j.stem.2024.03.002 PubMed PMID: 38579682; PubMed Central PMCID: PMC11212398.

97. Ferrari S, Jacob A, Beretta S, Unali G, Albano L, Vavassori V, et al. Efficient gene editing of human long-term hematopoietic stem cells validated by clonal tracking. Nat Biotechnol. 2020 Nov;38(11):1298–308. doi:10.1038/s41587-020-0551-y PubMed PMID: 32601433; PubMed Central PMCID: PMC7610558.

98. Wolff JH, Skov TW, Haslund D, Dorset SR, Revenfeld ALS, Aussel C, et al. Targeted gene editing and near-universal cDNA insertion of CYBA and CYBB as a treatment for chronic granulomatous disease. Nat Commun. 2025 Aug 12;16(1):7475. doi:10.1038/s41467-025-62738-2

99. Skov TW, Wolff JH, Haslund D, Revenfeld AL, van de Venn L, Dorset SR, et al. Treatment of GATA2 deficiency by allele-specific CRISPR-Cas9-directed gene correction in hematopoietic stem cells. Mol Ther. 2025 Nov 5;33(11):5644–60. doi:10.1016/j.ymthe.2025.07.038 PubMed PMID: 40739756; PubMed Central PMCID: PMC12628058.

100. Baik R, Cromer MK, Glenn SE, Vakulskas CA, Chmielewski KO, Dudek AM, et al. Transient inhibition of 53BP1 increases the frequency of targeted integration in human hematopoietic stem and progenitor cells. Nat Commun. 2024 Jan 2;15(1):111. doi:10.1038/s41467-023-43413-w

101. Schmidt M, Schwarzwaelder K, Bartholomae C, Zaoui K, Ball C, Pilz I, et al. High-resolution insertion-site analysis by linear amplification-mediated PCR (LAM-PCR). Nat Methods. 2007 Dec;4(12):1051–7. doi:10.1038/nmeth1103 PubMed PMID: 18049469.

102. Wang W, Bartholomae CC, Gabriel R, Deichmann A, Schmidt M. The LAM-PCR Method to Sequence LV Integration Sites. Methods Mol Biol. 2016;1448:107–20. doi:10.1007/978-1-4939-3753-0_9 PubMed PMID: 27317177.

103. Girard-Gagnepain A, Amirache F, Costa C, Lévy C, Frecha C, Fusil F, et al. Baboon envelope pseudotyped LVs outperform VSV-G-LVs for gene transfer into early-cytokine-stimulated and resting HSCs. Blood. 2014 Aug 21;124(8):1221–31. doi:10.1182/blood-2014-02-558163

104. Humbert O, Gisch DW, Wohlfahrt ME, Adams AB, Greenberg PD, Schmitt TM, et al. Development of Third-generation Cocal Envelope Producer Cell Lines for Robust Lentiviral Gene Transfer into Hematopoietic Stem Cells and T-cells. Mol Ther. 2016 Aug;24(7):1237–46. doi:10.1038/mt.2016.70 PubMed PMID: 27058824; PubMed Central PMCID: PMC5088759.

105. Leclerc D, Siroky MD, Miller SM. Next-generation biological vector platforms for in vivo delivery of genome editing agents. Curr Opin Biotechnol. 2024 Feb;85:103040. doi:10.1016/j.copbio.2023.103040 PubMed PMID: 38103518.

106. Haldrup J, Andersen S, Labial ARL, Wolff JH, Frandsen FP, Skov TW, et al. Engineered lentivirus-derived nanoparticles (LVNPs) for delivery of CRISPR/Cas ribonucleoprotein complexes supporting base editing, prime editing and in vivo gene modification. Nucleic Acids Res. 2023 Oct 13;51(18):10059–74. doi:10.1093/nar/gkad676 PubMed PMID: 37678882; PubMed Central PMCID: PMC10570023.

107. Chen K, Han H, Zhao S, Xu B, Yin B, Lawanprasert A, et al. Lung and liver editing by lipid nanoparticle delivery of a stable CRISPR-Cas9 ribonucleoprotein. Nat Biotechnol. 2025 Sep;43(9):1445–57. doi:10.1038/s41587-024-02437-3 PubMed PMID: 39415058; PubMed Central PMCID: PMC12000389.

108. Zhang Z, Baxter AE, Ren D, Qin K, Chen Z, Collins SM, et al. Efficient engineering of human and mouse primary cells using peptide-assisted genome editing. Nat Biotechnol. 2023 Apr 24. doi:10.1038/s41587-023-01756-1

109. Foss DV, Muldoon JJ, Nguyen DN, Carr D, Sahu SU, Hunsinger JM, et al. Peptide-mediated delivery of CRISPR enzymes for the efficient editing of primary human lymphocytes. Nat Biomed Eng. 2023 Apr 25;7(5):647–60. doi:10.1038/s41551-023-01032-2

110. Sahu SU, Castro M, Muldoon JJ, Asija K, Wyman SK, Krishnappa N, et al. Peptide-enabled ribonucleoprotein delivery for CRISPR engineering (PERC) in primary human immune cells and hematopoietic stem cells. Nat Protoc. 2025 Oct;20(10):2735–70. doi:10.1038/s41596-025-01154-8 PubMed PMID: 40032999.

111. Tommasi A, Cappabianca D, Bugel M, Gimse K, Lund-Peterson K, Shrestha H, et al. Efficient nonviral integration of large transgenes into human T cells using Cas9-CLIPT. Mol Ther Methods Clin Dev. 2025 Mar 13;33(1):101437. doi:10.1016/j.omtm.2025.101437 PubMed PMID: 40123742; PubMed Central PMCID: PMC11930092.

112. Roth TL, Puig-Saus C, Yu R, Shifrut E, Carnevale J, Li PJ, et al. Reprogramming human T cell function and specificity with non-viral genome targeting. Nature. 2018 Jul;559(7714):405–9. doi:10.1038/s41586-018-0326-5 PubMed PMID: 29995861; PubMed Central PMCID: PMC6239417.

113. Xie K, Starzyk J, Majumdar I, Rincones K, Tran T, Lee D, et al. Circular single-stranded DNA is a superior homology-directed repair donor template for efficient genome engineering [Internet]. Bioengineering; 2022 [cited 2026 May 3]. Available from: http://biorxiv.org/lookup/doi/10.1101/2022.12.01.518578 doi:10.1101/2022.12.01.518578

114. Letort G, Duclert A, Le Clerre D, Chion-Sotinel I, Salvatori R, Dessez E, et al. Circular single stranded DNA potentiates non-viral gene insertion in hematopoietic stem and progenitor cells. Nat Commun. 2025 Nov 19;16(1):10125. doi:10.1038/s41467-025-66318-2 PubMed PMID: 41257973; PubMed Central PMCID: PMC12630883.

115. Shy BR, Vykunta VS, Ha A, Talbot A, Roth TL, Nguyen DN, et al. High-yield genome engineering in primary cells using a hybrid ssDNA repair template and small-molecule cocktails. Nat Biotechnol. 2023 Apr;41(4):521–31. doi:10.1038/s41587-022-01418-8

116. Anzalone AV, Randolph PB, Davis JR, Sousa AA, Koblan LW, Levy JM, et al. Search-and-replace genome editing without double-strand breaks or donor DNA. Nature. 2019 Dec;576(7785):149–57. doi:10.1038/s41586-019-1711-4 PubMed PMID: 31634902; PubMed Central PMCID: PMC6907074.

117. Gelsinger DR, Vo PLH, Klompe SE, Ronda C, Wang HH, Sternberg SH. Bacterial genome engineering using CRISPR-associated transposases. Nat Protoc. 2024 Mar;19(3):752–90. doi:10.1038/s41596-023-00927-3 PubMed PMID: 38216671; PubMed Central PMCID: PMC11702153.

118. Yarnall MTN, Ioannidi EI, Schmitt-Ulms C, Krajeski RN, Lim J, Villiger L, et al. Drag-and-drop genome insertion of large sequences without double-strand DNA cleavage using CRISPR-directed integrases. Nat Biotechnol. 2023 Apr;41(4):500–12. doi:10.1038/s41587-022-01527-4 PubMed PMID: 36424489; PubMed Central PMCID: PMC10257351.

119. Pandey S, Gao XD, Krasnow NA, McElroy A, Tao YA, Duby JE, et al. Efficient site-specific integration of large genes in mammalian cells via continuously evolved recombinases and prime editing. Nat Biomed Eng. 2025 Jan;9(1):22–39. doi:10.1038/s41551-024-01227-1 PubMed PMID: 38858586; PubMed Central PMCID: PMC11754103.

120. Durrant MG, Perry NT, Pai JJ, Jangid AR, Athukoralage JS, Hiraizumi M, et al. Bridge RNAs direct programmable recombination of target and donor DNA. Nature. 2024 Jun;630(8018):984–93. doi:10.1038/s41586-024-07552-4 PubMed PMID: 38926615; PubMed Central PMCID: PMC11208160.

121. Ferreira da Silva J, Tou CJ, King EM, Eller ML, Rufino-Ramos D, Ma L, et al. Click editing enables programmable genome writing using DNA polymerases and HUH endonucleases. Nat Biotechnol. 2025 Jun;43(6):923–35. doi:10.1038/s41587-024-02324-x PubMed PMID: 39039307; PubMed Central PMCID: PMC11751136.

